# Overexpression of the transcription factor GROWTH-REGULATING FACTOR5 improves transformation of dicot and monocot species

**DOI:** 10.1101/2020.08.23.263947

**Authors:** Jixiang Kong, Susana Martín-Ortigosa, John Finer, Nuananong Orchard, Andika Gunadi, Lou Ann Batts, Dhiraj Thakare, Bradford Rush, Oliver Schmitz, Maarten Stuiver, Paula Olhoft, David Pacheco-Villalobos

## Abstract

Successful regeneration of genetically modified plants from cell culture is highly dependent on the species, genotype, and tissue-type being targeted for transformation. Studies in some plant species have shown that when expression is altered, some genes regulating developmental processes are capable of triggering plant regeneration in a variety of plant cells and tissue-types previously identified as being recalcitrant to regeneration. In the present research, we report that developmental genes encoding GROWTH-REGULATING FACTORS positively enhance regeneration and transformation in both monocot and dicot species. In sugar beet (*Beta vulgaris ssp. vulgaris*), ectopic expression of *Arabidopsis GRF5* (*AtGRF5*) in callus cells accelerates shoot formation and dramatically increases transformation efficiency. More importantly, overexpression of *AtGRF5* enables the production of stable transformants in recalcitrant sugar beet varieties. The introduction of *AtGRF5* and *GRF5* orthologs into canola (*Brassica napus* L.), soybean (*Glycine max* L.), and sunflower (*Helianthus annuus* L.) results in significant increases in genetic transformation of the explant tissue. A positive effect on proliferation of transgenic callus cells in canola was observed upon overexpression of *GRF5* genes and *AtGRF6* and *AtGRF9*. In soybean and sunflower, the overexpression of *GRF5* genes seems to increase the proliferation of transformed cells, promoting transgenic shoot formation. In addition, the transformation of two putative *AtGRF5* orthologs in maize (*Zea mays* L.) significantly boosts transformation efficiency and resulted in fully fertile transgenic plants. Overall, the results suggest that overexpression of *GRF* genes render cells and tissues more competent to regeneration across a wide variety of crop species and regeneration processes. This sets GRFs apart from other developmental regulators and, therefore, they can potentially be applied to improve transformation of monocot and dicot plant species.

## 2 Introduction

Plants have an impressive capability to regenerate new tissues, organs, or even an entire plant. External signals can trigger differentiated somatic cells to reprogram their developmental fate to repair wounded organs and to regenerate new tissues or whole plants, allowing plants to cope with environmental threats (Ikeuchi et al., 2016). This natural competence for regeneration is fundamental in plant propagation and tissue culture techniques that regenerate plants through *de novo* organogenesis or somatic embryogenesis using exogenously applied plant hormones (Ikeuchi et al., 2016; Kareem et al., 2016; Méndez-Hernández et al., 2019). The study of these regeneration pathways and their developmental regulators has enabled the establishment of protocols for the *in vitro* production and propagation of many plant species under controlled conditions (George et al., 2008; Sugiyama, 2015).

Tissue culture-induced regeneration is also important for transformation protocols in crop species. For example, monocot transformation methods predominantly depend on plant regeneration through somatic embryogenesis, whereas the regeneration of transgenic dicot crops is often achieved through organogenesis (Kausch et al., 2019). Regeneration potential of plant cells and organs varies widely among plant species, which often restricts transformation methods to a limited set of plant varieties or genotypes as well as target explants for DNA delivery (Cheng et al., 2004). The phenomenon of limited regenerative capability *in vitro*, better known as recalcitrance, makes the recovery of transgenic lines difficult or impossible in many plant species. This hampers the advance of fundamental research as well as the application of technologies relying on regeneration, including plant transformation and genome editing (Altpeter et al., 2016).

Traditionally, tissue culture-based regeneration methods are based upon the application of plant hormone combinations (especially auxins and cytokinins) on explants that are amenable to regeneration (Sugiyama, 2015). The establishment of a successful regeneration or transformation protocol often requires customization of the plant hormone ratios along with other tissue culture factors for each genotype, which is laborious and very often fails. Recently, several lines of experimental evidence suggest that inherent developmental programs are activated in response to tissue culture, which can reprogram the cell fate or activate undifferentiated cells in cultured explants (Ikeuchi et al., 2019; Sugimoto et al., 2019). Observations gained from model organisms such as *Arabidopsis* have shown that specific transcription factors can integrate the signals leading to cell reprogramming and the reacquisition of an embryonic or a meristematic fate (Gordon et al., 2007; Duclercq et al., 2011; Ikeuchi et al., 2013; Gaillochet and Lohmann, 2015; Kareem et al., 2015; Perez-Garcia and Moreno-Risueno, 2018). Such transcription factors are referred to as developmental regulators since they coordinate the spatial distribution of cells in an organized way during the development of organs and embryos.

Several genes encoding developmental regulators have been described to improve the regeneration efficiency in various plant species (Heidmann et al., 2011; Horstman et al., 2017; Gordon-Kamm et al., 2019). For example, the constitutive expression of *BABY BOOM* (*BBM*), a transcription factor of the AP2/ERF family with diverse functions in plant development, can promote cell proliferation and ectopic embryo formation in cotyledons and leaves of *Arabidopsis* (Boutilier et al., 2002). In a study aimed at elucidating the molecular mechanism of somatic embryogenesis, gain-of-function mutations in the gene coding for the shoot apical meristem identity regulator *WUSCHEL* (*WUS*) were found to induce embryo formation from various vegetative tissues in *Arabidopsis* (Zuo et al., 2002; Somssich et al., 2016). These initial observations led researchers to develop an approach using ectopic co-expression of *BBM* and *WUS* to greatly improve *in vitro* transformation in a variety of monocots, including several recalcitrant maize inbred lines, rice and sorghum (Lowe et al., 2016; Mookkan et al., 2017; Lowe et al., 2018). The positive influence of these genes on the regeneration in tissue culture was also reported in studies conducted in several dicot plant species, including highly recalcitrant crop varieties (Srinivasan et al., 2007; Arroyo-Herrera et al., 2008; Deng et al., 2009; Solís-Ramos et al., 2009; Heidmann et al., 2011; Bouchabke-Coussa et al., 2013; Florez et al., 2015).

Additional developmental genes have been proposed as candidates to further improve regeneration and transformation technologies in monocots and dicots (Gordon-Kamm et al., 2019; Kausch et al., 2019). For example, overexpression of an AP2/ERF transcription factor, WOUND INDUCED DEDIFFERENTIATION1 (WIND1) increased callus-induction and shoot regeneration in rapeseed, tomato, and tobacco (Iwase et al., 2015). WIND1 directly binds the promoter of another member of the AP2/ERF transcription factor family, *ENHANCER OF SHOOT REGENERATION1 (ESR1)* gene, whose overexpression similarly increases callus formation and shoot regeneration in *Arabidopsis* (Banno et al., 2001; Iwase et al., 2007). However, in all these reports the continued overexpression of developmental genes caused severe growth defects during further plant development, such as aberrant development of vegetative and reproductive organs and infertility. Therefore, use of these developmental genes requires restricting or eliminating their expression in the plant by methods such as gene excision or the use of inducible or tissue-specific promoters (Gallois et al., 2002; Lowe et al., 2016).

Here, we report that ectopic expression of a developmental regulator, *GROWTH-REGULATING FACTOR 5* (*GRF5*), has a positive effect in boosting regeneration and genetic transformation in various crop species using established organogenic or embryogenic regeneration systems. The *GRF* genes belong to a small plant-specific transcription factor family and they form a transcriptional complex with *GRF-INTERACTING FACTOR* (*GIF*) to regulate plant growth and development by providing cues to primordial cells of vegetative and reproductive organs (van der Knaap et al., 2000; Kim et al., 2003; Omidbakhshfard et al., 2015; Kim, 2019; Liebsch and Palatnik, 2020). Genetic analyses of plants with *AtGRF5* overexpression and loss-of-function mutations in *Arabidopsis* revealed that this transcription factor is involved in the regulation of the cell division and expansion during leaf development (Horiguchi et al., 2005; Horiguchi et al., 2006; Debernardi et al., 2014). In addition, *GRF5* overexpression in *Arabidopsis* increased chlorophyll content and chloroplast number per cell, and delayed leaf senescence (Vercruyssen et al., 2015). We observed that stable transformation of *GRF5* genes in sugar beet accelerated shoot organogenesis and resulted in significant improvements in genetic transformation, including recalcitrant varieties. Ectopic expression of *AtGRF5* and orthologs also resulted in significant increases in the formation and development of transgenic callus in canola and increases in the production of developing transgenic shoots in soybean and sunflower. In maize, transformation of *AtGRF5* orthologs showed a higher rate of embryogenic callus growth leading to an increased recovery of transgenic plants. Unlike other reported developmental regulators, no detrimental pleiotropic effects were observed in transgenic events overexpressing *GRF* genes in the tested crops, other than possible reduction in *de novo* root initiation on soybean shoots. The role of *GRF* genes to improve transformation protocols based on either shoot organogenesis or somatic embryogenesis is discussed.

## 3 Material and Methods

KWS SAAT SE & Co. KGaA and BASF will provide the material described here to academic groups under a material transfer agreement for non-commercial use in research.

### 3.1 *Agrobacterium* strains and binary plasmids

All bacterial strains and binary vectors used in this study were produced according to standard procedures (Russell and Sambrook, 2001). The *Agrobacterium* strains harboring the constructs used to transform sugar beet, canola, soybean, sunflower and maize, including the expression cassettes for the *GRF5* genes or the fluorescent control reporters, are described in the Supplementary Table 1. The plasmids used for the soybean immature embryo bombardment are also described in the Supplementary Table 1.

### 3.2 Sugar beet transformation

The multigerm inbred lines 9BS0448, 1RV6183, 7RV5706H and 8RV6921 were used in the sugar beet experiments. *Agrobacterium*-mediated transformation of sugar beet was performed by using the procedure and culture media as described elsewhere (Kishchenko et al., 2005) with minor modifications. The media composition is provided in the Supplementary Table 2. Briefly, to induce friable callus cultures, leaf explants from micropropagated shoots were incubated in callus induction medium at 30°C in the dark. The *Agrobacterium* culture was centrifuged and resuspended to an OD_600_ of 0.8 in liquid co-culture medium. Callus pieces were harvested and cultivated with *Agrobacterium* in co-culture medium solidified with 10 g/l agar for at least 2 days. Transformed calli were cultured in shoot regeneration medium for selection of transgenic shoots. Transgenic calli were transferred to fresh shoot regeneration medium every 3 weeks at least four times. Developing shoots were isolated and clonally propagated in shoot multiplication medium. Co-transformation experiments were performed by mixing *Agrobacterium* strains harboring the *AtGRF5* construct and the tdTomato control construct in a 1:1 volume ratio. To regenerate untransformed plants, callus cultures were subjected to the same procedure by using medium without aminoglycoside antibiotic. Transformation efficiency was estimated as the frequency of transgenic plants normalized by the number of starting explants.

### 3.3 Canola hypocotyl transformation

Canola transformation was performed on the genotype BNS3 based on the hypocotyl transformation method (Radke et al., 1988) with modifications. Media components used are detailed in Supplementary Table 3. Seedlings were prepared by surface sterilizing seeds in 70% ethanol and sowing on germination medium in PlantCon(tm) boxes and growing for 4 days in the dark at 23°C. *Agrobacterium rhizogenes* strain SHA001 was used for delivery of the gene cassettes into hypocotyl segments of *B. napus* seedlings (Mankin et al., 2007). Two experiments were conducted. The first experiment had 6 replicates over time each with 3 operators testing *AtGRF5, BnGRF5-LIKE*, or *DsRed* control vectors, and the second had three replicates over time each performed by three operators testing *AtGRF5, AtGRF6, AtGRF9* or *DsRed* control vectors. *A. rhizogenes* was grown to OD_600_ of 1.0 and subsequently diluted to 0.1 with liquid infection medium. Hypocotyl explants 7 to 10 mm in length were prepared from 4 day-old-seedling. After cutting, the explant was dipped in the *Agrobacterium* suspension and placed on filter paper on solid co-culture medium in 15 × 100 mm plates. Plates containing ∼50 explants were sealed with 3M Micropore(tm) tape and cultivated in 16 light/8 dark photoperiod at 23°C. After 3 days, all explants were transferred to recovery medium, sealed with tape, and cultivated for 7 days. Explants were then transferred to four rounds of selection, each cycle lasting 14 days. Shoots of at least 1 cm in size were removed to rooting medium. Explants were scored for callus growth and *DsRed* expression 38 days after transformation. Putative transformation efficiency was calculated as [(# events with either *DsRed* expression or rooting on selection/# explants transformed) *100].

### 3.4 Soybean primary-node transformation

Soybean transformation was performed using the cultivars, Jake and CD215, using the primary-node transformation method with modifications (Olhoft et al., 2007; Chang Y.-F., 2012). Seedlings were grown for donor material by plating sterile soybean seeds on solid germination medium (see Supplementary Table 4) in PlantCons(tm) and placed under light (150 μm^-2^s^-2^) at 26°C for 7 to 8 days under 18 light/6 dark photoperiod. *Agrobacterium rhizogenes* strain SHA017 was grown and resuspended to OD_600_ 1.5. Explants were prepared from seedlings by removing the roots, half of the hypocotyl, one cotyledon, and the epicotyl above the primary node. Approximately 50 explants were incubated in the *Agrobacterium* suspension for 30 minutes, then placed on co-cultivation medium and incubated at room temperature for 5 days. Explants were transferred to selection medium containing 3 μM imazapyr and cultivated at 26°C under light. After 3 weeks on selection, the explants were transferred to Oasis^®^ growing media and subsequently scored for *DsRed* fluorescence on developing shoots at the primary node. Explants were regularly watered with 5.7 µM IAA for 3 weeks. Elongated shoots were removed, placed on Oasis^®^ growing media, and watered with 5.7 µM IAA / 2 µM imazapyr for 7 days. The number of rooted plants was recorded, and a subset was sent to the GH for molecular characterization of T_0_ and T_1_ materials.

### 3.5 Sunflower shoot induction and *Agrobacterium*-mediated transformation

Seeds of sunflower (cv. RHA280) were harvested from plants grown under greenhouse conditions and stored at 4°C in the dark for up to 1 year. Sunflower cotyledon explants, *Agrobacterium* inoculation and sunflower transformation were conducted using the low inoculum with long co-culture (LI/LC) method (Zhang and Finer, 2016) at inoculum density of 10^2^ CFU/ml. After 2 weeks of co-culture on SIM solid medium, the explants were transferred onto solid shoot elongation medium (SEM). GFP expression in adventitious shoots was observed after 1 week on the SEM medium (3 weeks after inoculation). The *GRF5*s and GFP control vectors were transformed in triplicate (n ≥ 10).

### 3.6 Maize transformation

Transgenic plants from the maize inbred line A188 were produced according to the method described by (Ishida et al., 2007). Transformation efficiency was calculated as the frequency of inoculated immature embryos that regenerated at least one transgenic plant.

### 3.7 Microscopy

Sugar beet and maize *in vitro* cultures were photographed using a SteREO Discovery.V12 stereomicroscope (Carl Zeiss Microscopy GmbH, Jena, Germany). To quantify the embryogenic callus area in maize, 36 randomly taken pictures of calli obtained from independent embryos for each construct were used. The callus area was quantified in the pictures by using the open source software Fiji (Schindelin et al., 2012). DsRed fluorescence signal in canola and soybean primary node transformation experiments was monitored by using a Zeiss SV11 stereomicroscope coupled with the mercury vapor short-arc illuminator HBO 100W (Carl Zeiss Vision International GmbH, Aalen, Germany). The rhodamine filter was used for *DsRED2* (563 nm excitation max and 582 emission max) protein visualization. In soybean somatic embryogenesis-based transformation and sunflower transformation, tissue growth and GFP expression were monitored using a MZFLIII stereomicroscope (Leica, Heerbrugg, Switzerland) equipped with a “GFP-2” filter set (excitation 480 ± 40 nm; emission 510) and a pE-100 light-emitting diode (LED) lamp (Andover, Hampshire, UK) as an excitation light source.

### 3.8 Molecular analysis

*Sugar beet and maize:* Genomic DNA was extracted by standard procedures from leaf tissue and used for Multiplex TaqMan assays to measure the copy number of integrated T-DNA in transgenic sugar beet and maize plants. In both cases detection of endogenous genes (*BvEF1* of sugar beet and *ZmEF1* of maize) were combined with the detection of either *PAT* or *NPTII* gene, respectively, in a duplex qPCR. Primers and probes were designed using the Primer Express 3.0 software for Real-Time PCR (Applied Biosystems Foster City, California, USA), selecting the option MGB TaqMan probes with amplicon length 60 to 80 bp, and using the default parameters of the software. The oligonucleotide sequences used in TaqMan assays are provided in the Supplementary Table 5. qPCRs were carried out on an Applied Biosystems QuantStudio 3 and 5 Real-Time PCR Systems (Applied Biosystems, Foster City, California, USA) using 80 - 150 ng of genomic DNA as template, Maxima Probe qPCR Master Mix (2X), with ROX (ThermoFisher Scientific, Waltham, Massachusetts, USA), 250 nM MGB probe each and 900 nM primer each in a total volume of 10 μl. Amplification and quantification were performed with the following cycling conditions: 2 min 50°C, 5 min at 95°C for initial DNA denaturation, followed by an amplification program of 40 cycles at 95°C for 15 s and 60°C for 30 sec. The fluorescence threshold was manually set above the background level. Negative controls without DNA template and verified control samples with 1 copy and 2 copies of the T-DNA for either *PAT* or *NPTII* assay were run in parallel and used for absolute quantification and copy number determination.

*Soybean*: Genomic DNA was extracted from leaf tissue using a modified protocol for Wizard(R) Magnetic 96 DNA Plant System from Promega (Madison, Wisconsin, USA). The copy number of integrated T-DNA in transgenic soybean plants was measured in duplex TaqMan assays. Detection of endogenous 1-copy genes (*GmLectin*) was combined with the detection of *AHAS* gene in a duplex qPCR. Primers and probes were designed using the Primer Express 3.0.1 software for Real-Time PCR (Applied Biosystems, Foster City, California, USA), selecting the option TaqMan^®^ Quantification probes, and using the default parameters of the software. The oligonucleotide sequences used in TaqMan assays are provided in the Supplementary Table 5. qPCRs were carried out on an Applied Biosystems QuantStudio 7 Real-Time PCR Systems (Applied Biosystems, Foster City, California, USA) using isolated genomic DNA as template, Perfecta^®^ qPCR ToughMix^®^ Low ROX(tm) PCR master mix (QuantaBio, 2x), 200 nM Taqman probe and 900 nM forward and reverse primers in 5 μl. Amplification and quantification were performed with the following cycling conditions: 2 min 95°C for initial DNA denaturation, followed by an amplification program of 40 cycles at 95°C for 2 min and 60°C for 1 min. The fluorescence threshold was manually set above the background level and verified control samples with 1 copy of the T-DNA for *AHAS* assay were run in parallel and used for absolute quantification and copy number determination.

### 3.9 qRT-PCR analysis

Total RNA from leaf samples was isolated with InviTrap^®^ Spin Plant RNA Mini Kit (INVITEK Molecular, Berlin, Germany) and reverse transcribed using Invitrogen(tm) Oligo (dT) primers (Fisher Scientific GmbH, Schwerte, Germany) and Invitrogen(tm) SuperScript(tm) III Reverse Transcriptase (Fisher Scientific GmbH, Schwerte, Germany). 20 ng of reverse transcribed RNA was used as template in the real-time PCR using Applied Biosystems(tm) QuantStudio(tm) 6 Flex (Fisher Scientific GmbH, Schwerte, Germany). Specific primers binding the 3’-UTR of the NOS terminator were used for the detection of the transgene transcripts. The qPCR reactions for *AtGRF5* and the endogenous reference (*Beta vulgaris* subsp. *vulgaris CASEIN KINASE 1-LIKE PROTEIN 11*) were performed in technical duplicates by using the following oligonucleotides: NOSt-F, 5’-AAC GTC ATG CAT TAC ATG TTA-3’; NOSt-R, 5’-CGG TCT TGC GAT GAT TAT CA-3’; S989-F, 5’-GAG GAA CTA GAC ATG GGG ATA CAT-3’; S990-R, 5’-GCG ATA CAA AGT AGA CAT TAG AAC TC-3’. ΔΔCt values were calculated and transcript levels given in log2 scale relative to the transgenic T_0_ event (L50) displaying the lowest *AtGRF5* overexpression.

### 3.10 Data analysis

In all experimental data sets, statistical significance of differences in average was determined with a Welch’s t-test.

### 3.11 Identification of *GRF5* homologs and sequence alignment

To identify the ortholog genes of *AtGRF5* from sugar beet, canola, soybean, sunflower and maize, AtGRF5 amino acid sequence was used as the query to perform BLAST against respective plant-specific protein sequence databases (proteins from most actual annotated genome assemblies from NCBI and EnsemblPlants). Initial candidates were selected based on the BLAST results, and further analyzed by global, multiple alignments: true redundancy (= identical protein sequences) and outliers (= suspicious sequences, as revealed by manually checking using multiple alignments) were removed first. Then, the initial candidates were ranked based on true pairwise global identities by comparing with AtGRF5 (program NEEDLE from EMBOSS): ranking was based on global identity, thus also the less-conserved C-terminal region of the transcription factors is considered here. Based on this ranking, top candidates were selected, and confirmed to align best (global identity) with AtGRF5 when compared with all 9 *Arabidopsis* GRF-family members in global alignments (back search). A summary of sequence identities found according to the blastp searches is shown in Supplementary Table 6.

GRF5 protein sequences were aligned using CLC DNA Workbench 8.0 (Qiagen) with the following settings: gap open cost, 10; gap extension cost, 1; end gap cost, as any other; alignment, very accurate. The conserved QLQ and WRC domains in the GRF crop orthologs were identified by using the *Arabidopsis* consensus amino acid sequences (Kim and Tsukaya, 2015) (Supplementary Figure 3).

### 3.12 Accession numbers

Sequence data from this article can be found in the GenBank/EMBL libraries under accession numbers AY102638 (*AtGRF5*, At3g13960), XP_010666262 (*BvGRF5-LIKE*), XM_022719744.1 (*BnGRF5-LIKE*), XP_006589198.1 (*GmGRF5-LIKE)*, XM_022128855.1 (*HaGRF5-LIKE*), GRMZM2G105335 (*ZmGRF5-LIKE1*), and GRMZM5G893117 (*ZmGRF5-LIKE2*).

## 4 Results

### 4.1 Overexpression of *GRF5* promotes shoot organogenesis and increases transformation efficiency in sugar beet

During an *Agrobacterium*-mediated transformation experiment in sugar beet line 9BS0448, we noticed that a large number of shoots was unexpectedly regenerated from transgenic calli expressing the *AtGRF5* gene under the CaMV 35S promoter (*2×35S*::*AtGRF5*). That observation prompted us to hypothesize that overexpression of *AtGRF5* in callus cells positively influences shoot organogenesis. To test, we transformed leaf-derived callus from the same line with *Agrobacterium* containing *2×35S*::*AtGRF5* and compared to the control construct *2×35S*::*tdT*, which expresses the tdTomato fluorescent protein. Compared to the control experiment, the calli transformed with the *AtGRF5* transgene showed enhanced and rapid shoot formation in all experiments, confirming our initial observation (Figure 1A). The number of regenerated shoots was significantly greater from callus expressing *AtGRF5* at the end of the second (S2), third (S3) and fourth (S4) round of selection compared to the control (Figure 1B). Interestingly, an average of 30 shoots was already formed from *AtGRF5*-transformed callus at the S2 round in each experiment, whereas in the control experiment, shoot formation was only visible at the selection round S3 (Figure 1B). Next, qPCR analyses confirmed that 97.6% of the regenerated shoots were transgenic in all experiments. Strikingly, callus transformed with *2×35S*::*AtGRF5* produced more transgenic events per experiment than the *2×35S*::*tdT* control cultures, resulting in a significant 6-fold increase on transformation efficiency (Figure 1C). Similarly, transformation efficiency increased when sugar beet calli were inoculated with a mixture of *Agrobacterium* cultures containing either the *2×35S*::*AtGRF5* or the *2×35S*::*tdT* construct. Transgenic plants containing both constructs were regenerated, resulting in an averaged co-transformation efficiency of 19.6% (Supplementary Figure 1). Compared to the experiments performed with the *tdTomato* construct alone, transgenic plants containing the *2×35S*::*tdT* cassette were more efficiently produced in the co-transformation experiments, suggesting that *AtGRF5* overexpression improves the recovery of transgenic events containing a construct of interest (Supplementary Figure 1). In addition, leaf explants from four out of five randomly selected T_0_ events produced callus cultures with enhanced shoot formation capacity compared to non-transgenic callus cultures (Figure 2). This correlated with the expression level of *AtGRF5* of the lines (Figure 2). Sugar beet plants that were transformed with *2×35S*::*AtGRF5* grew well *in vitro* and showed no obvious differences in their phenotype compared with the transgenic control plants growing under the same conditions (Supplementary Figure 2). Collectively, our results indicate that the ectopic expression of *AtGRF5* boosts the switch from callus phase to organogenesis and therefore, efficiently improves the transformation in sugar beet.

**Figure 1.**
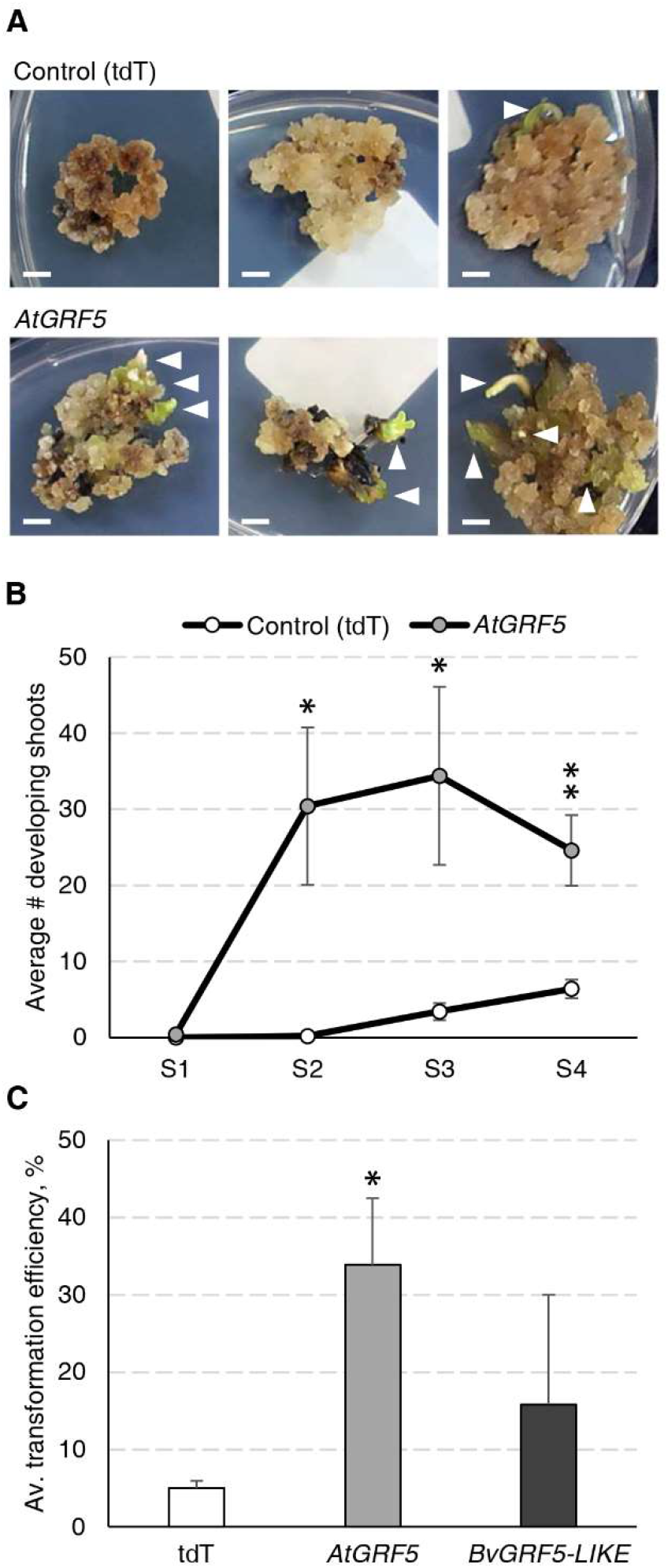
Detailed analysis of the *AtGRF5* overexpression in sugar beet transformation experiments. Transformations with a construct expressing the tdTomato (tdT) were done in parallel as a control. **(A)** Representative pictures of transformed callus on selection medium. White arrow heads indicate developing shoots. Scale bar = 0.5 cm. **(B)** Number of shoots regenerated across four rounds of culture on selection medium (S1 to S4). The mean values ± SEM from 5 biological replicates for the tested constructs are represented. **(C)** Average transformation efficiency of the experiments performed with the construct to overexpress *AtGRF5* or the putative sugar beet ortholog *BvGRF5-LIKE* construct and compared to the control construct. The transformation efficiency values are indicated as the mean ± SEM from at least 3 biological replicates per tested construct. In both graphs, differences as compared to the control are not significant unless indicated otherwise; error bars indicate standard error; * = p<0.05; ** = p<0.01.

**Figure 2.**
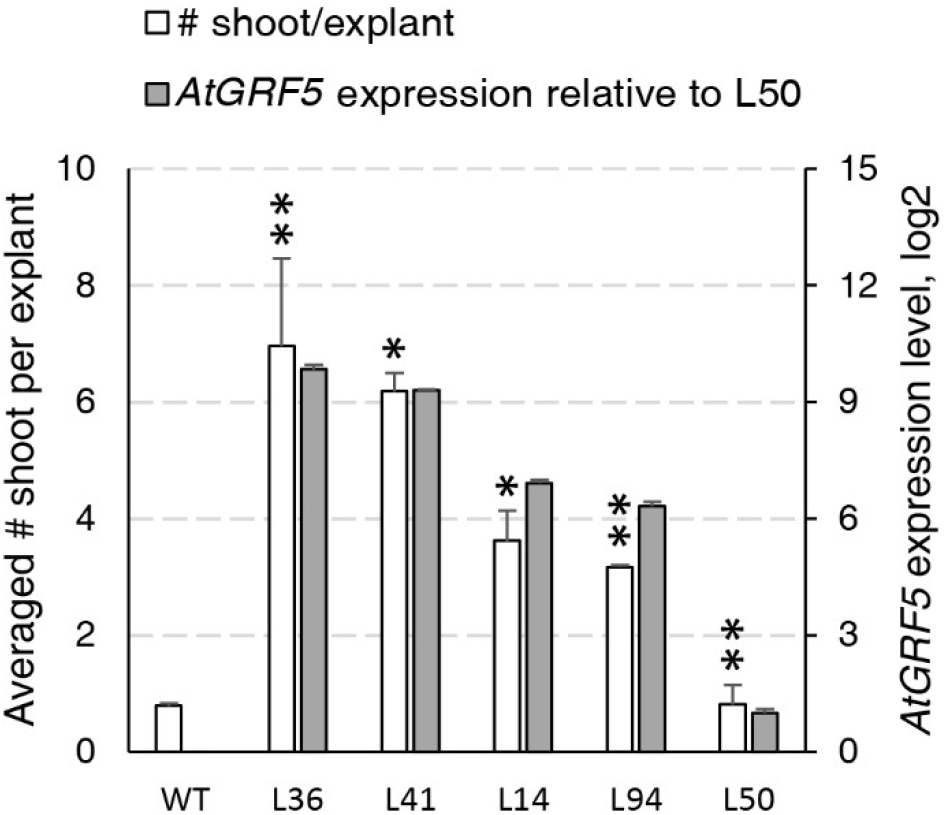
Characterization of the regeneration capacity in sugar beet T0 events expressing *AtGRF5*. Shoot regeneration capacity of friable callus cultures produced from five independent transgenic events containing the *2×35S*::*AtGRF5* construct (L36, L41, L14, L94 and L50) was compared with a non-transgenic plant regenerated from callus (WT) as a control (white bars). *AtGRF5* expression level is represented with gray bars. The expression values were normalized to the low expressing event L50. The gene expression values are indicated as the mean ± SEM from 2 technical replicates, whereas the shoot/explant values are indicated as the mean ± SEM from at least 50 explants isolated from each transgenic event. Differences as compared to the control are not significant unless indicated otherwise; error bars indicate standard error; * = p<0.05; ** = p<0.01.

We suspected that the sugar beet *GRF5* homolog might also influence shoot organogenesis. The putative ortholog in sugar beet was identified by blastp searches and named *BvGRF5-LIKE* (Supplementary Figure 3 and Supplementary Table 6). To test, we followed the same experimental design and transformed sugar beet callus of the line 9BS0448 with a construct to overexpress the *BvGRF5-LIKE* and compared with the control *tdTomato*. The transformation of *BvGRF5-LIKE* in sugar beet callus resulted in an increased transformation efficiency, although not statistically significant (Figure 1C).

### 4.2 *GRF5* promotes shoot organogenesis in different sugar beet varieties, including recalcitrant ones

It is well known that sugar beet transformation is highly genotype dependent (Gurel et al., 2008). Consequently, we investigated the effect of *AtGRF5* in the transformation efficiency of three other sugar beet varieties selected from the KWS germplasm. Leaf-derived callus obtained from the three inbred lines 1RV6183, 7RV5706H and 8RV6921, with differing degrees of recalcitrance, was transformed with either the *2×35S*::*AtGRF5* construct or the *2×35S*::*tdT* construct, respectively. The three lines showed enhanced transformability when the *AtGRF5* construct was used (Figure 3A). The line 1RV6183 displayed transformation efficiencies with the control construct (6.2%) and the *2×35S*::*AtGRF5* construct (27.8%) that are comparable to the 9BS0448 variety (Figure 3A and Figure 1C). The lines 7RV5706H and 8RV6921 showed an average transformation efficiency with the control construct of 1.8% and 0%, respectively, indicating a higher level of recalcitrance to transformation of these lines compared to 1RV6183 and 9BS0448 (Figure 3A and Figure 1C). By contrast, the introduction of the *2×35S*::*AtGRF5* construct into the calli resulted in transformation efficiencies higher than the control experiment for the 7RV5706H (20.7%) and 8RV6921 (6.2%) sugar beet varieties (Figure 3A). Interestingly, all shoots that were regenerated from the recalcitrant 8RV6921 line contained the *AtGRF5* gene, while the transformation of the *2×35S*::*tdT* construct did not produce any shoot (Figure 3A and 3B). Overall, these results suggest that the overexpression of *AtGRF5* can overcome recalcitrance to transformation in sugar beet.

**Figure 3.**
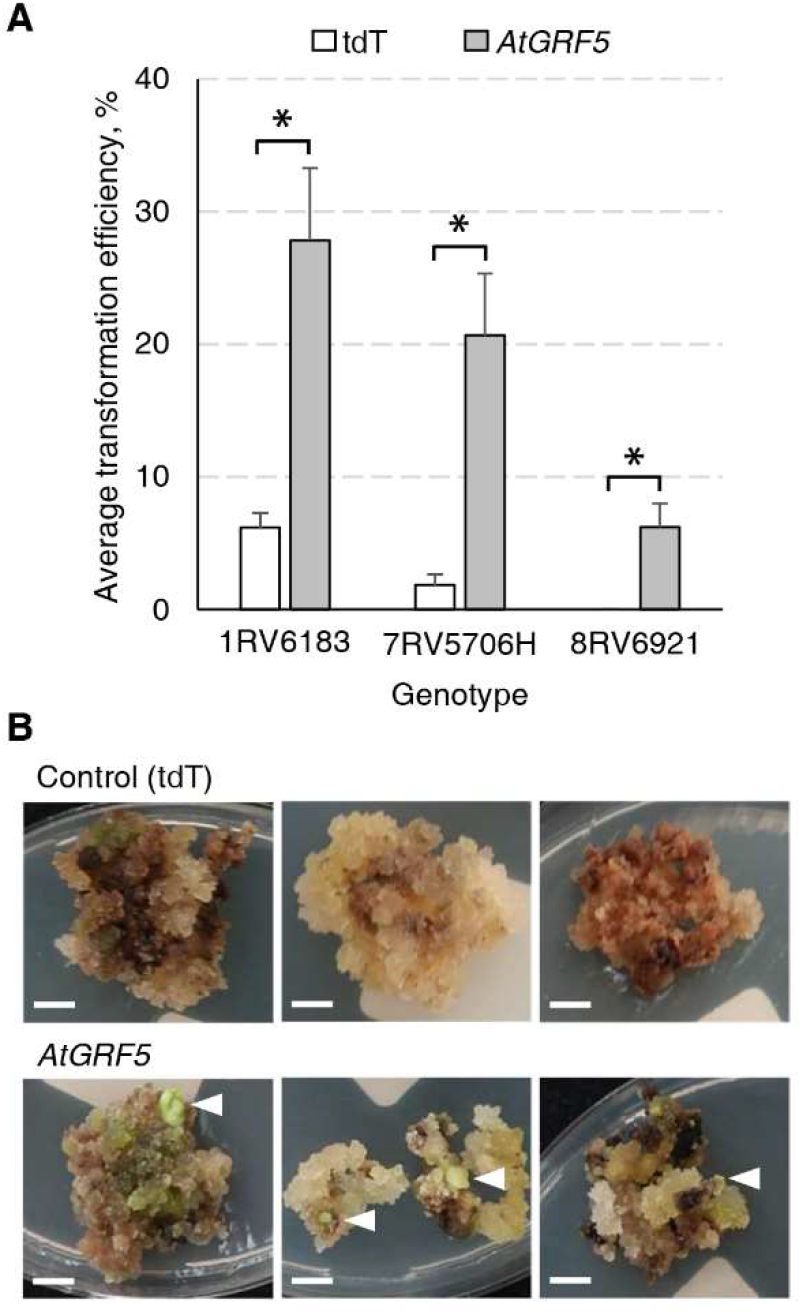
Overexpression of *AtGRF5* enables transformation of recalcitrant sugar beet varieties. Transformations done with the tdTomato (tdT) and the *AtGRF5* constructs were compared. **(A)** Average transformation efficiency of 3 sugar beet genotypes. Across the tested genotypes, the level of recalcitrance to shoot regeneration increases from left to right, the genotype 8RV6921 being the most recalcitrant. The transformation efficiency values are indicated as the mean ± SEM from at least 3 biological replicates per construct and genotype. Differences as compared to the control are not significant unless indicated otherwise; error bars indicate standard error; * = p<0.05. **(B)** Representative pictures of transformed calluses of the recalcitrant 8RV6921 genotype in selection medium. White arrow heads indicate developing shoots. Scale bar = 0.5 cm.

### 4.3 Overexpression of *GRF5, GRF6*, and *GRF9* increases transgenic callus production in canola

Next, we explored whether ectopic expression of *GRF* orthologs and paralogs could result in increased transgenic callus and shoot formation in canola. In addition to *AtGRF5* and its canola ortholog *BnGRF5-LIKE*, two further members of the AtGRF transcription factor family, i. e. *AtGRF6* and *AtGRF9*, were also selected in order to test whether positive effects on transformation are specific to GRF5 or can be more generally attributed to the GRF gene family. The rationale to test the aforementioned two candidate genes was based on sequence homology with *AtGRF6* being the most similar and *AtGRF9* being the most distant GRF paralog compared to *AtGRF5*. In addition, *AtGRF5* and *AtGRF9* are both strongly expressed in the lower half of 6-day old leaf primordia while *AtGRF6* is expressed mainly near the midvein in *Arabidopsis* (Horiguchi et al., 2005).

The GRF ortholog and paralog expression cassettes were introduced into the canola genome using a hypocotyl transformation method (Radke et al., 1988). Although shoot production occurs through organogenesis in this method, the shoot primordia arise from callus tissue induced from hypocotyl segments that are targeted for T-DNA integration. To test whether overexpression of *GRF* genes could increase transgenic callus or shoot production, explants from cultivar BNS3 were transformed with *AtGRF5, BnGRF5-LIKE, AtGRF6, AtGRF9*, or *DsRed* control vectors in two experiments. Overexpression of all the *GRF5* orthologs and paralogs resulted in significant increases in explants with *DsRed* fluorescing sectors than compared to the control vector (Figure 4A). At this step in the process, the hypocotyl has prominent callus formation, especially at the ends of the hypocotyls. To better understand if the *DsRed* fluorescence was localized to developing callus tissues, *DsRed* expression was further characterized in one experiment. The percent of explants with *DsRed* callus was found to be significantly greater on explants transformed with *AtGRF5* (*x* = 53.3; SE = 3.4; *p* = 1.1 * 10^−4^) or *BnGRF5-LIKE* (x = 66.7; SE = 4.0; *p* = 2.7 * 10^−6^) compared to *DsRed* control (*x* = 28.7; SE = 3.5), indicating that the majority of *DsRed* sectors were in callus tissues. *DsRed* fluorescence was also notably stronger in those explants with callus overexpressing *AtGRF5* or *BnGRF5-LIKE* (Figure 4B). The high *DsRed* expression suggests the callus is undergoing rapid cell proliferation compared to the control. Shoots that formed were transplanted onto rooting media and scored for rooting or *DsRed* fluorescence on the plantlet. Although there was a significant increase in *DsRed* expressing callus, no significant difference was found in the number of rooted and/or *DsRed* expressing shoots between *GRF5* transformed events compared to *DsRed* control events. These results could be interpreted as either *GRF5* overexpression in canola was unfavorable for shoot formation or the high *DsRed* expression is inhibiting further plant development.

**Figure 4.**
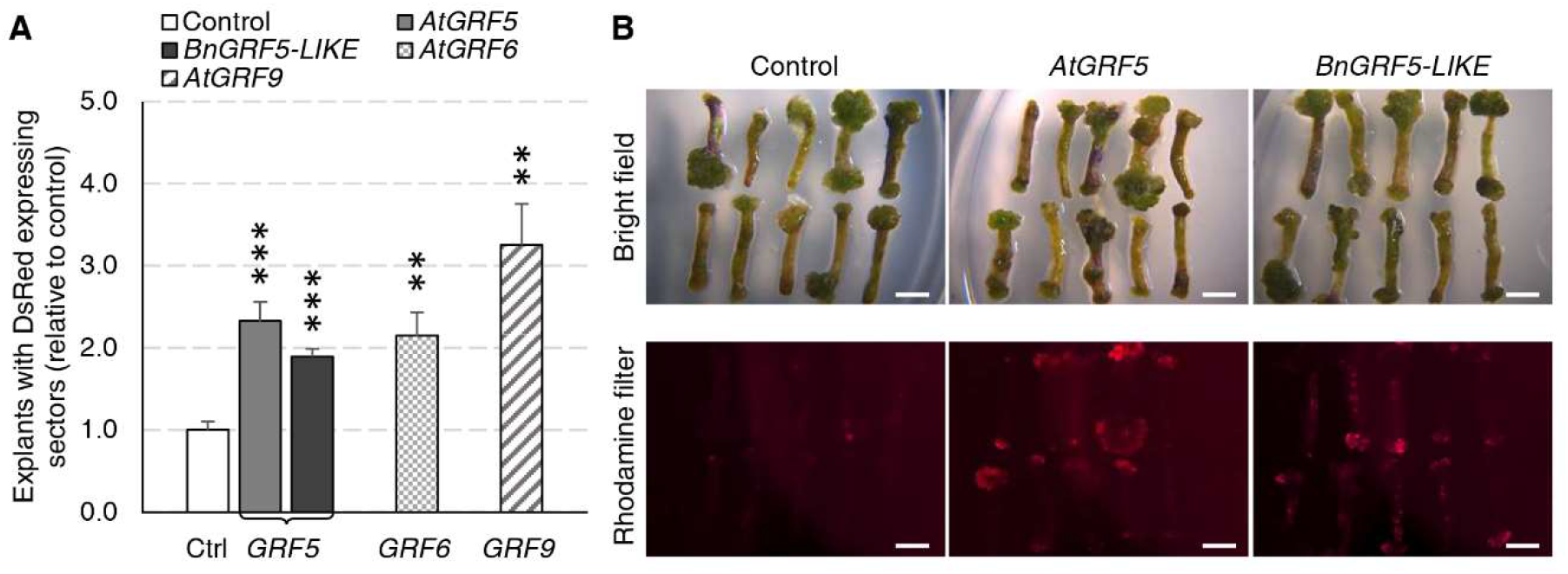
Canola hypocotyl sections producing callus increased in explants transformed with *AtGRF5, BnGRF5-LIKE, AtGRF6*, and *AtGRF9* relative to *DsRed* control vectors. **(A)** Explants with DsRed expressing sectors (relative to control) were analyzed in two experiments where the first experiment had 6 biological replicates over time testing *AtGRF5, BnGRF5-LIKE*, or *DsRed* control vectors, and the second had three biological replicates over time testing *AtGRF5, AtGRF6, AtGRF9* or *DsRed* control vectors. Explants overexpressing *AtGRF5 or BnGRF5-LIKE* resulted in a significant increase in DsRed sectors 38 days after transformation relative to *DsRed* control vector. In all graphs, differences as compared to the control are not significant unless indicated otherwise; error bars indicate standard error; ** = p<0.01, *** = p<0.001. **(B)** Representative pictures of explants 21 days after transformation transformed with vectors containing *DsRed* control, *AtGRF5*, and *BnGRF5-LIKE* show a relative increase in size and intensity of DsRed in overexpression cassettes of both *GRF5* orthologs compared to *DsRed* control. Callus development on the hypocotyl segments under white light is shown on the top panel and DsRed fluorescence under a rhodamine filter on explants in the lower panel. Scale bar=5mm.

### 4.4 Overexpression of *GRF5* promotes shoot regeneration directly from axillary meristems via organogenesis in soybean

We first wanted to address whether overexpression of *GRF5* orthologs from *Arabidopsis* and soybean can increase transgenic shoot production in soybean directly from axillary meristematic cells. To answer this question, a series of experiments were initiated using two cultivars, Jake and CD215, and assayed for the presence of *DsRed* fluorescence in shoots developing at the seedling’s primary node 27 days after transformation. In the first experiment, prepared explants of the cultivar Jake were transformed with the *DsRed* control, *AtGRF5*, or *GmGRF5-LIKE*. In three experiments, explant material from the cultivar CD215 was transformed with *GmGRF5-LIKE, AtGRF5*, or *DsRed* control vectors. Significantly more explants developed shoots expressing *DsRed* when transformed with *AtGRF5* or *GmGRF5-LIKE* compared to the *DsRed* control vector for both cultivars (Figure 5A). In addition, the explants transformed with *GRF5* genes, especially *GmGRF5-LIKE*, frequently had more shoot development per explant giving it a swollen appearance at the primary node. Interestingly, *DsRed* fluorescence was also notably stronger in shoots developing from *GRF5* transformed tissue compared to the *DsRed* control (Supplementary Figure 4A). The appearance of more meristem initials on the primary node suggests that overexpression of *GRF5* is increasing meristem and shoot formation at the axillary nodes.

**Figure 5.**
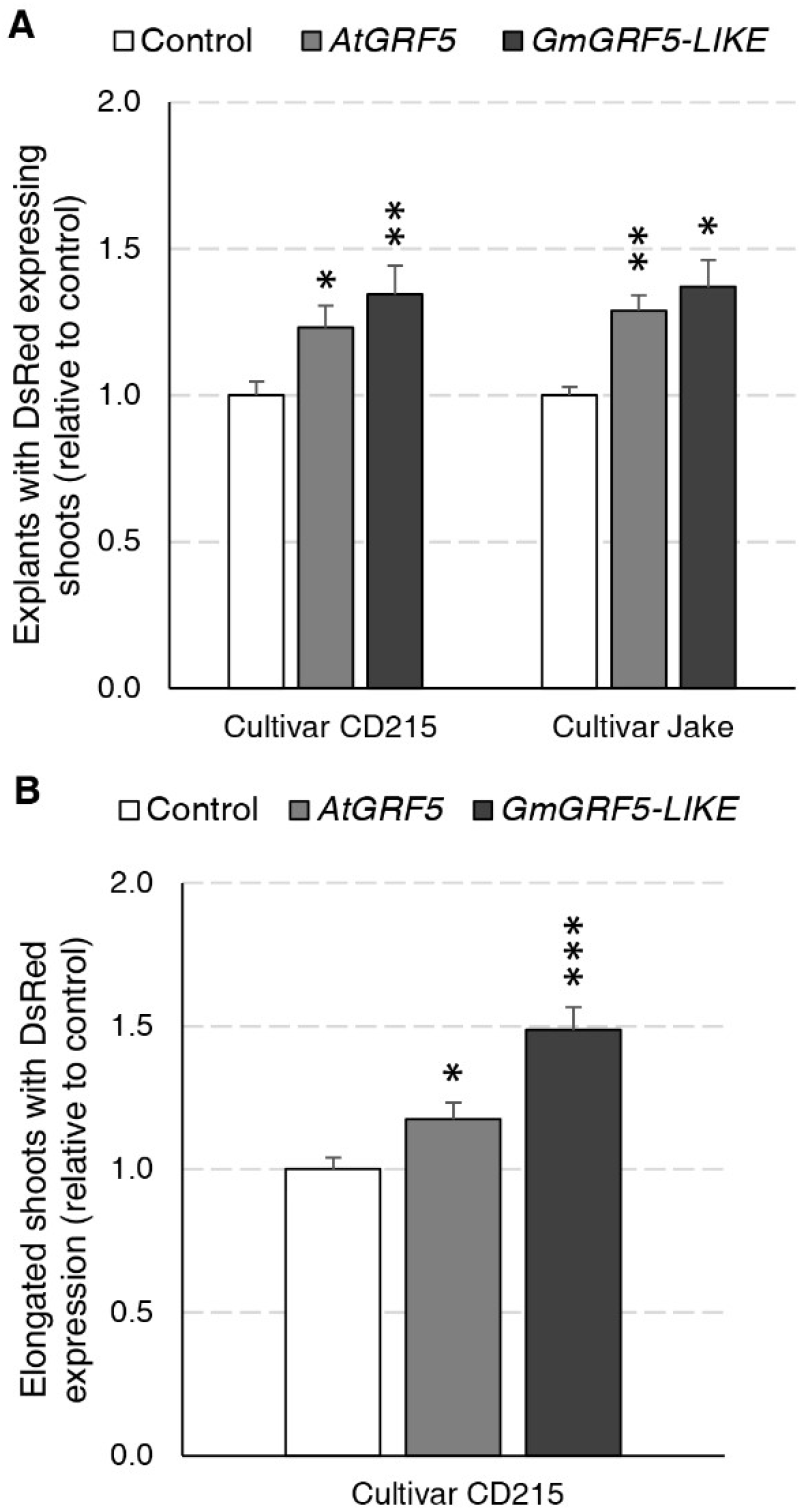
Soybean regeneration potential is increased in two cultivars expressing *AtGRF5* or *GmGRF5-LIKE* transcription factors relative to control. Explants of cultivar CD215 were transformed across three experiments each with three biological replicates testing *AtGRF5* or *DsRed* control vectors, *GmGRF5-LIKE* or *DsRed* control vectors, and *GmGRF5-LIKE, AtGRF5*, or *DsRed* control vectors. In a fourth experiment, prepared explants of the cultivar Jake were transformed with the *DsRed* control, *AtGRF5*, or *GmGRF5-LIKE* across four biological replicates. **(A)** Percent of explants with DsRed expressing shoots (normalized to control) significantly increased in explants transformed with either *GRF5* ortholog in both cultivars tested 27 days after transformation. **(B)** For CD215 experiments, elongated shoots were removed, rooted on imazapyr selection, and scored for DsRed expression over a 5-week period. Overexpression of *AtGRF5* and *GmGRF5-LIKE* resulted in the production of more DsRed expressing and/or rooted shoots. In both graphs, differences as compared to the control are not significant unless indicated otherwise; error bars indicate standard error; * = p<0.05; ** = p<0.01, *** = p<0.001.

Given that more explants produced shoots expressing *DsRed* with *GRF5* overexpression, we wanted to know if the increase in shoot production would also result in an increase of stable, transgenic plants. The experiments described above were continued until shoots elongated to 10-15 cm in length (Supplementary Figure 4B). Afterwards, one elongated shoot per explant was removed, placed on selection media for seven days and then scored for the presence or absence of roots. Despite the increase in shoot production on explants overexpressing *GRF5* 27 days after transformation, the observed differences in transgenic plant production between *GRF5* and *DsRed* control experiments for both cultivars were not statistically significant due to high experimental variability. In CD215, an average of 51.3% and 51.2% of explants produced rooted shoots for *AtGRF5* and *GmGRF5-LIKE*, respectively, compared to 44.7% of the control explants. Similarly, in Jake, an average of 50.2% of explants produced rooted shoots when transformed with *AtGRF5*, 47.2% with *GmGRF5-LIKE*, and 37.1% with *DsRed* control. To understand if the shoots expressing *DsRed* did not elongate or if the elongated shoots did not root, we scored all elongated shoots for *DsRed* expression after one week on rooting (Figure 5B). When compared to the control, explants transformed with *GRF5* genes produced significantly more elongated shoots expressing *DsRed* in the CD215 cultivar (Figure 5B). These results suggest that *GRF5* genes increased production of elongated, transgenic shoots, and reduced root formation negatively impacted the whole plant regeneration.

To confirm stable T-DNA integration in the T_0_ plant, copy number analysis of the *AtAhasL* gene in 126 independent events across all replicates was performed for CD215. The results confirmed that the transgene was indeed present in 99% of the rooted plants (Supplementary Table 7). Because overexpression of developmental genes has been associated with abnormal plant development and sterility, we continued to grow 42 events of CD215 until the R6 stage in the greenhouse. We observed no unusual morphological abnormalities in the developing plants and seeds of any of the events. Transgene inheritance into the T_1_ generation was then determined by a presence/absence assay for the *AtAhasL* gene in immature embryos using TaqMan (Supplementary Table 8). In all events, the *AtAhasL* gene was detected in the T_1_ generation in at least one progeny. A chi-square test of independence was performed on single copy T_0_ events to examine the relation between the genes, *GmGRF5-LIKE, AtGRF5*, and control cassette (*DsRed*) and the presence of the *AtAhasL* gene in the T_1_ generation. There was no significant relationship between the constructs and the ability to transmit the *AtAhasL* gene to the progeny, *X*^*2*^ (2, *N* = 405) = 5.2, *p* = 0.08, suggesting normal inheritance of the *AtGRF5* and *GmGRF5* genes. In several cases, the number of null segregants was greater than expected, which could be explained by small sample size and/or the occurrence of a chimeric T_0_ plant, which is common using organogenesis.

### 4.5 Overexpression of *GRF5* promotes transgenic shoot production from cotyledons via organogenesis in sunflower

Regeneration of transgenic sunflower plants remains problematic. Therefore, we wanted to test if overexpression of *GRF5* could increase transgenic shoot production or regeneration of plants using a low inoculum/low coculture transformation method that targets sunflower cotyledons (Zhang and Finer, 2016). Although shoot induction directly from the cotyledons via organogenesis is high in sunflower, transformed cells are usually concentrated at the cut sides where shoot formation rarely occurs (Zhang and Finer, 2016). Mature cotyledons of the cultivar, RHA280, were prepared and transformed with overexpression cassettes of *AtGRF5*, a putative sunflower ortholog, *HaGRF5-LIKE* (Supplementary Figure 3), or a GFP control vector. Explants that were transformed with *HaGRF5-LIKE* produced significantly more (*p*=0.005) GFP expressing shoots per explant than the control vector, while the number of explants transformed with *AtGRF5* was two-fold greater, albeit not statistically significant (Figure 6A). In addition, GFP expressing sectors were more expansive, not only on the explants, but also on the shoots when compared to the control (Figure 6B).

**Figure 6.**
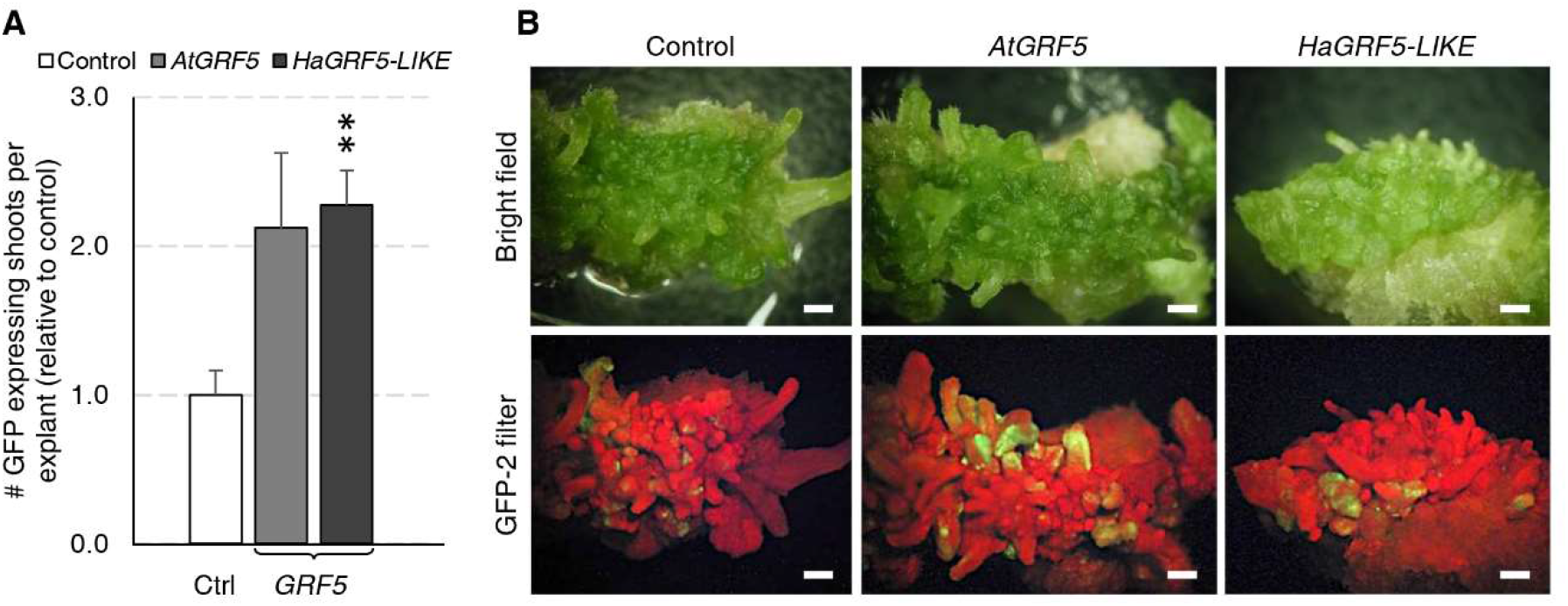
Overexpression of *AtGRF5* and the ortholog *HaGRF5-LIKE* in sunflower (cv. RHA280) has a positive effect on the formation of GFP expressing shoots per explant. A Welch’s t-test was performed to determine significance of differences in averages from three experiments with 7 biological replicates for GFP control, 6 biological replicates for *AtGRF5*, and 4 biological replicates for *HaGRF5-LIKE* vectors. **(A)** The number of GFP expressing shoots was significantly higher in *HaGRF5-LIKE* overexpressed explants relative to control, while a positive trend was observed on explants transformed with *AtGRF5* (*p = 0*.*079*). Differences as compared to the control are not significant unless indicated otherwise; error bars indicate standard error; ** = p<0.01. **(B)** The size and frequency of GFP expressing shoots is more pronounced in explants transformed with *GRF5*. Representative pictures of explants with GFP expression under bright field (top) and GFP fluorescence (bottom) at 3 weeks post *Agrobacterium* inoculation with either *GFP* control, *AtGRF5, HaGRF5-LIKE* vectors are shown. Scale bar = 1 mm.

### 4.6 Overexpression of *GRF5* improves somatic embryogenesis-based transformation in maize

To explore whether the *GRF5* gene could boost the regeneration of transgenic plants through somatic embryogenesis, we employed a transformation protocol to introduce GRF5 constructs in the maize inbred line A188 (Ishida et al., 2007). To this end, the putative homologs *ZmGRF5-LIKE1* and *ZmGRF5-LIKE2* were identified as previously described for other crops (Supplementary Figure 3) and cloned under the *BdEF1* promoter to ensure high expression level. Binary plasmids containing the expression cassette for either *ZmGRF5-LIKE1, ZmGRF5-LIKE2, AtGRF5* and *tdTomato* were transformed into immature embryos. Over 50% of the embryos transformed with any of the two *ZmGRF5-LIKE* homologs developed type I embryogenic callus that were bigger than the control experiment at the end of the second selection round on callus induction medium (Supplementary Figure 5). Surprisingly, the size of the embryogenic calli in the transformation with the *AtGRF5* construct remained comparable to the control at the same time point (Supplementary Figure 5). Quantitative analysis of callus area corroborated that the transformation of *ZmGRF5-LIKE* genes significantly increased the average embryogenic callus area compared to the control, whereas the transformation of the *Arabidopsis* counterpart had a very limited effect on the callus area (Figure 7A). These observations point out that, unlike *AtGRF5*, the maize *GRF5* genes might intensify the proliferation of scutellum-derived embryogenic callus. In contrast to the control and the *AtGRF5* transformations, a large amount of proliferating calli was obtained from the embryos transformed with *ZmGRF5* at the end of the callus selection rounds (Figure 7B). Next, qPCR analysis confirmed the presence of the T-DNA in most regenerated maize plants. In general, the experiments done with the *ZmGRF5-LIKE1* constructs showed a higher average transformation efficiency than the control (Figure 7C). Although a similar efficiency was observed when the *ZmGRF5-LIKE2* was transformed, the effect of this ortholog was not statistically significant from controls due to a greater experimental variation. On the other hand, the transformation of the *AtGRF5* gene in maize did not improve the efficiency of transgenic plant recovery (Figure 7C). The enhanced transformation efficiency is consistent with the increased embryogenic callus growth and revealed that the overexpression of *ZmGRF5-LIKE* genes improves transformation in maize.

**Figure 7.**
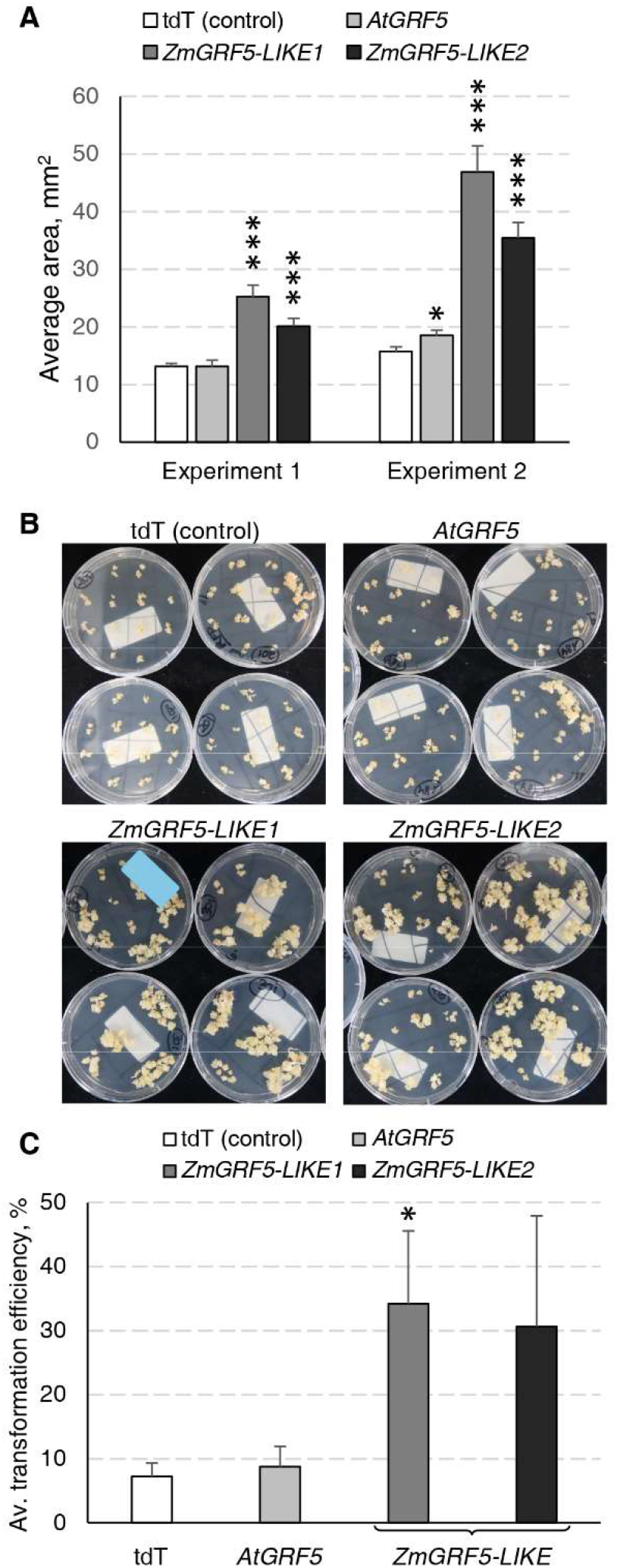
Analysis of the *GRF5* genes in maize transformation experiments. In all experiments, constructs for either *tdTomato* (tdT), *AtGRF5, ZmGRF5-LIKE1* or *ZmGRF5-LIKE2* overexpression were used. **(A)** Quantification of the area of developing embryogenic calluses at 39 days after *Agrobacterium* inoculation. The area values are indicated as the mean ± SEM from 36 randomly selected calluses that were obtained from transformed immature embryos in each experiment. **(B)** Representative transgenic callus cultures at the end of the callus selection phase, before transferring to medium for regeneration of transformed plants. **(C)** Average transformation efficiency of the experiments performed in maize with the *GRF5* and the tdT (control) constructs. The transformation efficiency values are indicated as the mean ± SEM from at least 3 biological replicates for *tdTomato, AtGRF5* and *ZmGRF5-LIKE1*; and 2 biological replicates for *ZmGRF5-LIKE2*. In the graphs, differences as compared to the control are not significant unless indicated otherwise; error bars indicate standard error; * = p<0.05; *** = p<0.001.

Five randomly selected maize T_0_ plants for each *ZmGRF5-LIKE* construct were grown in the greenhouse until maturity and compared with plants transformed with the *tdTomato* control construct. All cultivated T_0_ maize plants showed normal organ morphology and produced a similar average number of seeds (Supplementary Figure 6 and 7). In addition, segregation analysis showed mendelian inheritance of the transgenes in all tested T_1_ progenies (Supplementary Table 9). In summary, these results demonstrate that the overexpression of *ZmGRF5-LIKE* genes does not impair the growth and fertility in maize and the transgene can be transmitted to the next generation.

## 5 Discussion

Despite decades of research on plant tissue culture and genetic engineering, the ability to efficiently regenerate fertile plants from transformed somatic tissues and cells is still limited. Even though there has been considerable progress enabling *in vitro* organogenesis and somatic embryogenesis in a variety of crops, transformation is often restricted to very few crop varieties, limiting the application of genome editing and transformation to crop improvement (Altpeter et al., 2016). Expression of developmental genes to boost regeneration has been reported as promising strategy to overcome this hurdle in transformation of plants of agricultural interest (Nagle et al., 2018; Gordon-Kamm et al., 2019). Here, we have discovered that the overexpression of the transcription factor *GRF5* from *Arabidopsis* and/or its homologs increases transformation efficiency in sugar beet and maize. In addition, overexpression of *GRF5* homologs improves transgenic shoot formation in soybean and sunflower, and transgenic callus cell proliferation in canola.

### 5.1 *GRF5* overexpression boosts transgenic plant production by promoting shoot organogenesis in sugar beet

Creation of transgenic plants by shoot organogenesis from friable callus has been reported in sugar beet (Gurel et al., 2008). In these transformation protocols, transgenic callus cells were easily obtained, but the regeneration of transgenic plantlets was time consuming, difficult, and ultimately resulted in low transformation frequencies (Dhalluin et al., 1992; Kishchenko et al., 2005). In our study, we first discovered that the overexpression of *AtGRF5* in sugar beet friable callus greatly promoted and accelerated shoot development, increasing the efficiency of transgenic plant production and, at the same time, decreasing its turnaround time (Figure 1). According to previous research, the recalcitrance in sugar beet is likely due to the low number of cells morphologically competent for regeneration, which are difficult to access for transformation as they are immersed within a large number of noncompetent cells within the callus (Gurel and Wren, 1995; Joersbo, 2007). The high transgenic shoot formation rates suggest that sugar beet callus cells overexpressing *AtGRF5* might have acquired a regenerative advantage over the large population of noncompetent cells, which is in consonance with the function of *GRF* genes in regulating cell division and the pluripotent competence of cells and tissues in model plant species (Horiguchi et al., 2005; Lee et al., 2018). During *de novo* shoot organogenesis, cell division is required for both the generation of a new cell mass and the acquisition of shoot regeneration competency (Ikeuchi et al., 2019). In a genetic study, the *grf1/2/3/4* quadruple *Arabidopsis* mutant was shown to produce a significant portion of seedlings lacking shoot apical meristem (Kim and Lee, 2006), indicating that *GRF*-mediated cell proliferation might regulate shoot meristem establishment. Thus, *AtGRF5* overexpression might confer competence to the callus cells for *de novo* organogenesis facilitating the development of transgenic shoots. Further supporting that idea, friable callus obtained from stably *AtGRF5*-transformed leaf explants displayed enhanced *de novo* shoot formation compared to wild-type callus cultures in our experiments, which correlated with the *AtGRF5* expression level (Figure 2).

More recently, a protocol for producing transgenic sugar beet plants from callus cells was published after 14 years of continuous optimization of tissue culture parameters whereby a 10% transformation efficiency was achieved (Kagami et al., 2015). By contrast, the overexpression of *AtGRF5* led to transformation frequencies remarkably higher than 10%. Furthermore, the *AtGRF5* overexpression also enabled efficient recovery of co-transformed events, being the first time that co-transformation is reported in this crop. Interestingly, the callus transformed with *BvGRF5-LIKE* showed more variable frequencies of transgenic plant production compared to the *AtGRF5*-expressing callus, as illustrated by the large standard error of the mean transformation efficiency. That could be attributed to a differential response of the transformed callus cells to the overexpression of *AtGRF5* and *BvGRF5-LIKE* and their respective regulation or mode of action in sugar beet.

On the other hand, several reports have shown that sugar beet breeding lines and cultivars generally exhibit diverse *in vitro* responses to plant regeneration from friable callus, which illustrates strong genotype dependency of the regeneration in that crop (Dovzhenko and Koop, 2003; Ivic-Haymes and Smigocki, 2005; Tomita et al., 2013). Importantly, in our analysis *AtGRF5* overexpression dramatically improved the recovery of transgenic events in several sugar beet inbred lines, including a genotype that was not amenable for transformation up to date (Figure 3). That demonstrates the potential of *GRF* genes in broadening the range of varieties that can be transformed in recalcitrant crops, such as sugar beet.

### 5.2 Overexpression of *GRF5* genes promotes transgenic callus proliferation in canola

Next, we wondered whether *GRF* overexpression could improve indirect shoot organogenesis in other crops. Therefore, we decided to investigate the response of canola hypocotyl explants to the overexpression of *AtGRF6* and *AtGRF9*, and compare with *AtGRF5* and its putative *Brassica napus* ortholog *BnGRF5-LIKE*. In our analysis, the overexpression of *AtGRF5* and *BnGRF5-LIKE* resulted in higher frequencies of callus with transgenic sectors. Likewise, *AtGRF6* and *AtGRF9* overexpression showed similar effects, indicating that *GRF* genes in general have a positive effect on promoting transgenic callus proliferation (Figure 4). Considering that, unlike several other members of the GRF family, *AtGRF9* has been shown to have an opposite function in regulating cell proliferation and organ growth (Omidbakhshfard et al., 2018), it is very intriguing that the overexpression of this *Arabidopsis* homolog also promoted proliferation of transgenic callus cells in canola. Unlike sugar beet, there was no improvement in transgenic plant production in canola with any of the tested *GRF* genes. This could be attributed to differences in the transformation method where the hypocotyl explant is directly targeted for T-DNA delivery and callus is initiated from the explant. Shoot formation from the callus only occurs after several rounds of subcultures, which may explain some of the differences in plant production between canola and sugar beet.

### 5.3 *GRF5* overexpression improves transgenic shoot formation in soybean and sunflower

To further explore the effect of GRF overexpression in organogenesis, *AtGRF5* and their respective soybean and sunflower orthologs were transformed using published methods, which are based on direct shoot organogenesis (Olhoft et al., 2007; Chang Y.-F., 2012; Zhang and Finer, 2016). *GRF5* overexpression in soybean resulted in visibly more meristematic initials at the axillary meristem, indicating increased cell division activity. Furthermore, transgenic shoot production at the axillary node was notably improved (Figure 5). Although most of the *GRF5*-expressing shoots developed well, a proportion of the elongating shoots that were expressing *DsRed* did not form roots, indicating that *GRF5* or *DsRed* expression may negatively affect adventitious root initiation. By contrast, overexpression of *GRF5* genes in sugar beet and canola plants did not impair root growth, suggesting that the effect of *GRF5* on rooting might be crop specific, dependent on the promoter used, or due to the combination of *GRF5* overexpression and the regeneration method. Similarly, the frequency of explants containing transgenic shoots significantly increased when *GRF5* genes were transformed in sunflower, suggesting an effect on proliferation of transformed tissues (Figure 6). The results obtained in these two crops are again consistent with the notion that *GRF* genes positively regulate cell division in growing plant organs (Kim, 2019). Indeed, *Arabidopsis* plants overexpressing *AtGRF5* develop enlarged leaves containing more cells, which was also accompanied with increased expression of cell cycle genes (Horiguchi et al., 2005; Gonzalez et al., 2010; Debernardi et al., 2014).

Overall, although *GRF* overexpression improved transgenic cell proliferation in canola, sunflower and soybean, transformation efficiency was not always increased. A possibility that cannot be ruled out is that differences in the transformation protocols, such as the target explant and phytohormones used for regeneration, might interact with the overexpression of *GRFs*. For instance, it was shown that the overexpression of *AtGRF5* increases the sensitivity of *Arabidopsis* leaves to cytokinins and stimulated leaf growth when co-expressed with cytokinin catabolic enzymes (Vercruyssen et al., 2011; Vercruyssen et al., 2015). Several research groups also link the *GRFs* with other hormone pathways including brassinosteroids, gibberellins and auxin (Che et al., 2016; Gao et al., 2016; Lee et al., 2018; Zhang et al., 2018). It remains to be determined how GRF is linked to the hormone pathways related to plant regeneration.

### 5.4 Overexpression of *ZmGRF5* genes enhances transformation based on somatic embryogenesis in maize

Besides the positive effect on organogenesis, further analyses uncovered that overexpression of *GRF5* genes promotes regeneration in somatic embryogenesis-based transformation protocols in maize. Qualitative and quantitative analysis revealed that the overexpression of *ZmGRF5-LIKE* genes enhanced the growth of embryogenic callus, suggesting that *ZmGRF5-LIKE* homologs might have a positive influence on cell proliferation, thereby boosting the formation of somatic embryos and enhancing the transformation efficiency (Figure 7). According to the nomenclature used by RefSeq and adopted by Debernardi et al. (2020) the *ZmGRF5* genes tested in our study are orthologous to the wheat *GRF1* gene, which has also showed a positive effect on wheat transformation in a concurrent research (Debernardi et al., 2020). Like in dicot plants, GRF-promoting cell division and organ growth has also been described in monocots (Nelissen et al., 2015), which is again in accordance with the enhanced growth of transgenic embryogenic callus reported here. Interestingly, unlike the *ZmGRF5-LIKE* genes, overexpression of *AtGRF5* in maize only slightly promoted proliferation of maize embryogenic callus and did not enhanced transformation efficiency, suggesting functional diversification of the monocot and dicot *GRF* genes.

Although little is known about the role of GRF genes in somatic embryogenesis, a recent publication demonstrated that the overexpression of *AtGRF1* resulted in increased sensitivity of Arabidopsis explants to 2,4-D treatment and improved induction of somatic embryogenesis (Szczygiel-Sommer and Gaj, 2019). In our maize transformation protocol, the induction of somatic embryogenesis is achieved on 2,4-D-containing medium. Therefore, one can hypothesize that the overexpression of *ZmGRF5-LIKE1* or *ZmGRF5-LIKE2* might increase the sensitivity of the maize explants to 2,4-D, consequently improving their embryogenic response.

Finally, a technological advance employing *Agrobacterium*-mediated introduction of the BABYBOOM (BBM) and WUSCHEL (WUS) transcription factors greatly improved transformation efficiency in monocot crops, including maize recalcitrant varieties (Lowe et al., 2016; Lowe et al., 2018). However, constitutive expression of these transcription factors led to impaired plant development and sterility, which obliged researchers to develop strategies to limit expression to the first steps of the transformation process (Lowe et al., 2016). By contrast, the transgenic plants overexpressing either *AtGRF5* or *GRF5* orthologs in this study showed normal growth until maturity and produced viable seeds.

### 5.5 Concluding remarks and future prospects

In summary, we have demonstrated that members of the GRF transcription factor family can be used to improve transformation protocols based on either organogenesis or embryogenesis. In light of our results, it appears that the GRF-mediated regeneration improves genetic transformation in various plant species, including dicotyledonous and monocotyledonous crops. Interestingly, high transformation efficiencies were achieved in recalcitrant sugar beet genotypes, revealing, that *GRF* genes can be applied to decrease genotype-dependency of transformation protocols.

This work presents the first evidence that members of the *GRF* gene family can trigger cell reprogramming to accomplish more efficient *in vitro* regeneration and transformation. We hypothesize that the effect of *AtGRF5* and its counterparts on transformation in the crops analyzed in this study, can be mostly attributed to the well-known role of GRF transcription factors in regulating crucial developmental processes, such as cell proliferation. Accumulating evidence determined that the growth of numerous plant organs is controlled by a conserved gene network consisting of *GRFs, GRF-INTERACTING FACTOR* transcriptional co-regulators (*GIF*) and miR396 (Kim and Kende, 2004; Liebsch and Palatnik, 2020). Among other GRFs, AtGRF5 was also shown to interact with AtGIF1 (Horiguchi et al., 2005; Vercruyssen et al., 2014). Preliminary data suggest that co-expression of *AtGIF1* and *AtGRF5* induces a more pronounced effect on regeneration (data not shown). In line with this finding, researchers at UC Davis concurrently observed that overexpression of a chimeric protein containing GRF and its transcriptional co-activator GIF greatly increased the transformation efficiency in wheat, rice and citrus (Debernardi et al., 2020). In addition to the interaction with GRFs, GIF1 was shown to physically interact with SWI/SNF chromatin remodeling members to regulate transcription of various genes (Debernardi et al., 2014; Nelissen et al., 2015). The SWI/SNF complex is known to be antagonistic to the polycomb PRC2 complex (Wilson et al., 2011). Interestingly, mutations in the PRC2 complex subunits led to upregulation of downstream targets *WIND3* and *LEC2*, which promote regeneration (Ikeuchi et al., 2015). One can hypothesize that GRF5 might interact with the GIF1-SWI/SNF complex in order to inhibit the polycomb PRC2 complex, thereby releasing its repression on other developmental regulators, consequently triggering cell reprogramming (Ikeuchi et al., 2015). Continuous efforts to unravel the function of *GRF* genes in the context of transcriptional modulation of key meristem or embryonic regulators would contribute to our fundamental understanding of the molecular pathways determining cell reprogramming during plant *in vitro* regeneration. In addition, further optimization of protocols would help to find specific conditions in which GRF-mediated regeneration boosting leads to higher transformation efficiencies in recalcitrant crops.

## Supporting information

Kong et al (2020) supplementary information

## 7 Acknowledgments

We would like to thank Linda Schwerdtfeger, Franziska Mense, Heike Lehmann, and Robin Pfeil for excellent technical assistance on transformation experiments at KWS; Benjamin Gruber for critical reading of the manuscript; Florian Schauwecker for conducting sequence analyses to derive *GRF5* orthologs; James Silva Garcia for statistical analyses; Allan Wenck for critical review of manuscript; Fernando Ferreira for vector design and construction; Zhong Qi Wang, Shyam Barampuram, Harvinder Kaur, Minesh Patel, Yan Liu, Brandi Chappell and Mao Wang for executing and delivering on transformation experiments at BASF.

## 8 Author Contributions

Conceived and designed the experiments: JK, SMO, DP-V, PO, JF, OS, and MS. Performed the experiments: JK, SMO, NO, AG, JF, LB, DT, BR, and PO. Analyzed the data: OS and DP-V. Wrote the paper: JK, SMO, MS, OS, PO, and DP-V.

