## Supplementary material for "Overexpression of the transcription factor GROWTH-REGULATING FACTOR5 improves transformation of dicot and monocot species": Kong et al (2020) supplementary information

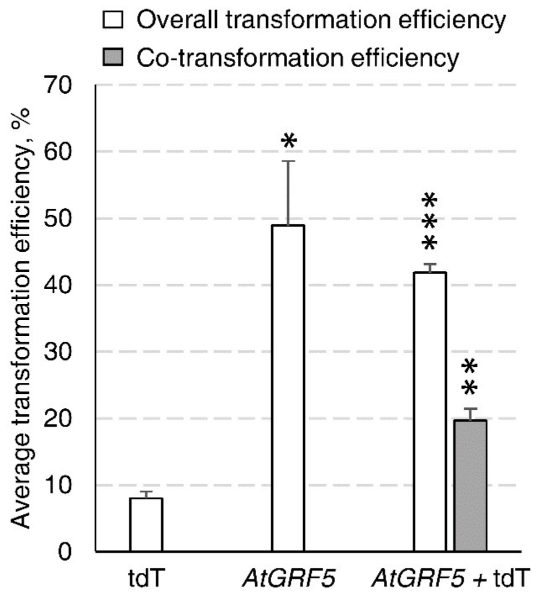


**Supplementary Figure 1**. *AtGRF5* enables co-transformation in sugar beet. Single transformations were performed with an *Agrobacterium*, harboring either the control *2x35S*::*tdT* construct (tdT) or the *2x35S*::*AtGRF5* construct, and co-transformation was performed with a 1:1 mixture of these two *Agrobacterium* strains (*AtGRF5* + tdT). The transformation efficiency values are indicated as the mean ± SEM from at least 3 biological replicates. Co-transformation efficiency was calculated with the events showing the presence of both *AtGRF5* and *tdTomato* cassettes. Differences as compared to the control (tdT, single transformation) are not significant unless indicated otherwise; error bars indicate standard error; * = $p<0.05$; ** = $p<0.01$; *** = $p<0.001$.

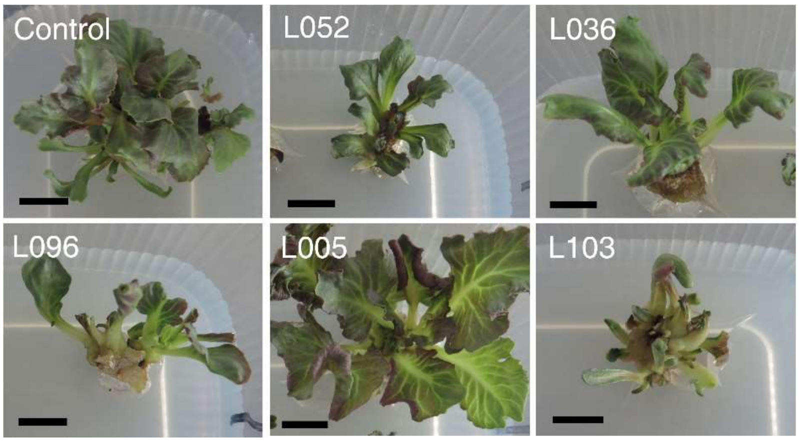


**Supplementary Figure 2**. Representative pictures of regenerated $T_0$ shoots growing *in vitro*. Transgenic shoots produced with the *2x35S::tdT* construct are indicated as control. L052, L036, L096, L005 and L103 correspond to independent transgenic shoots produced with the *2x35S*::*AtGRF5* construct. Scale bar = 1 cm.

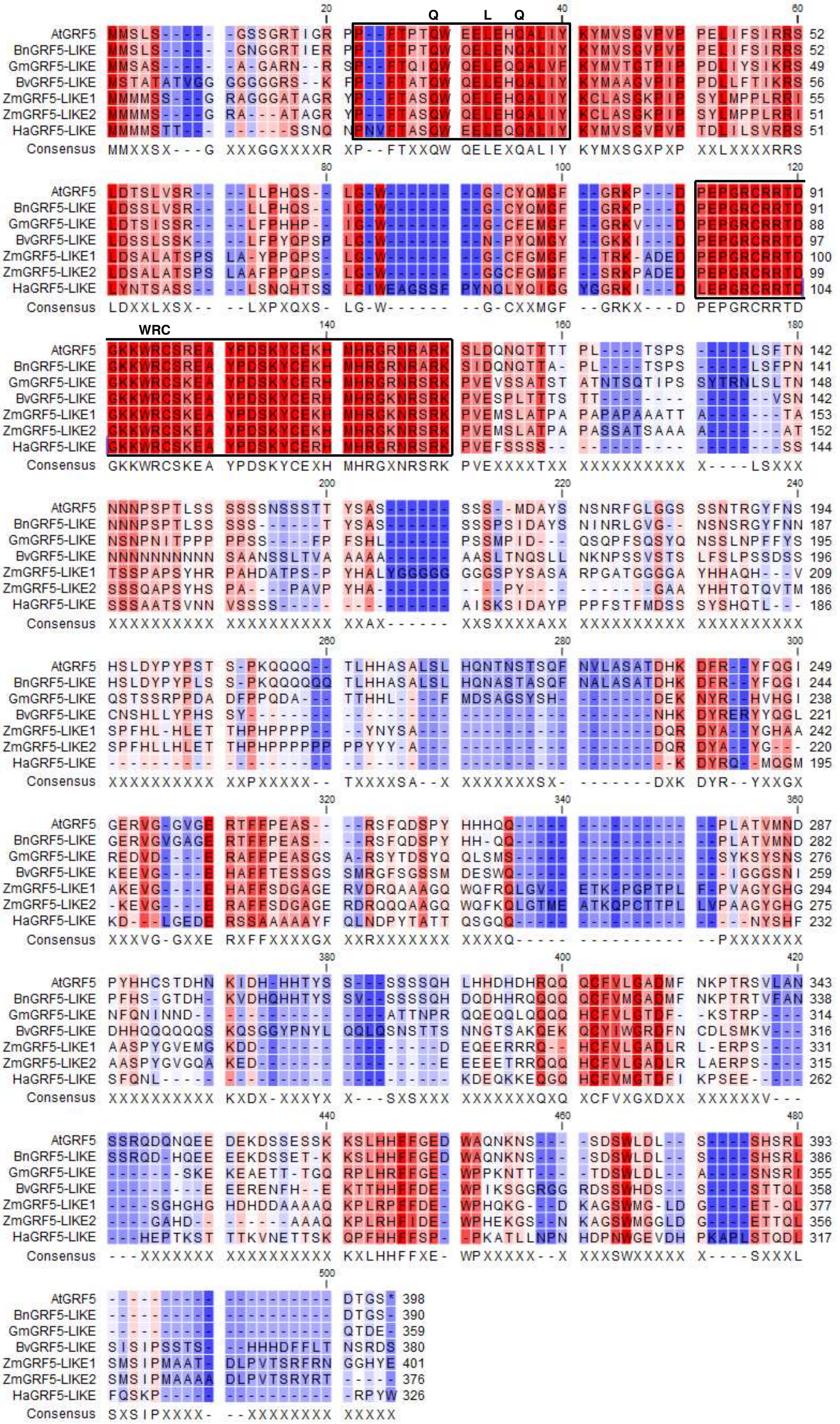


**Supplementary Figure 3**. Alignment of the protein sequence encoded by *AtGRF5* gene and representative *GRF5-LIKE* genes from *Brassica napus* (*Bn*), *Glycine max* (*Gm*), *Zea mays* (*Zm*), *Beta vulgaris* (*Bv*) and *Helianthus annuus* (*Ha*). Conserved QLQ and WRC domain are shown in boxes . *At*: *Arabidopsis thaliana*

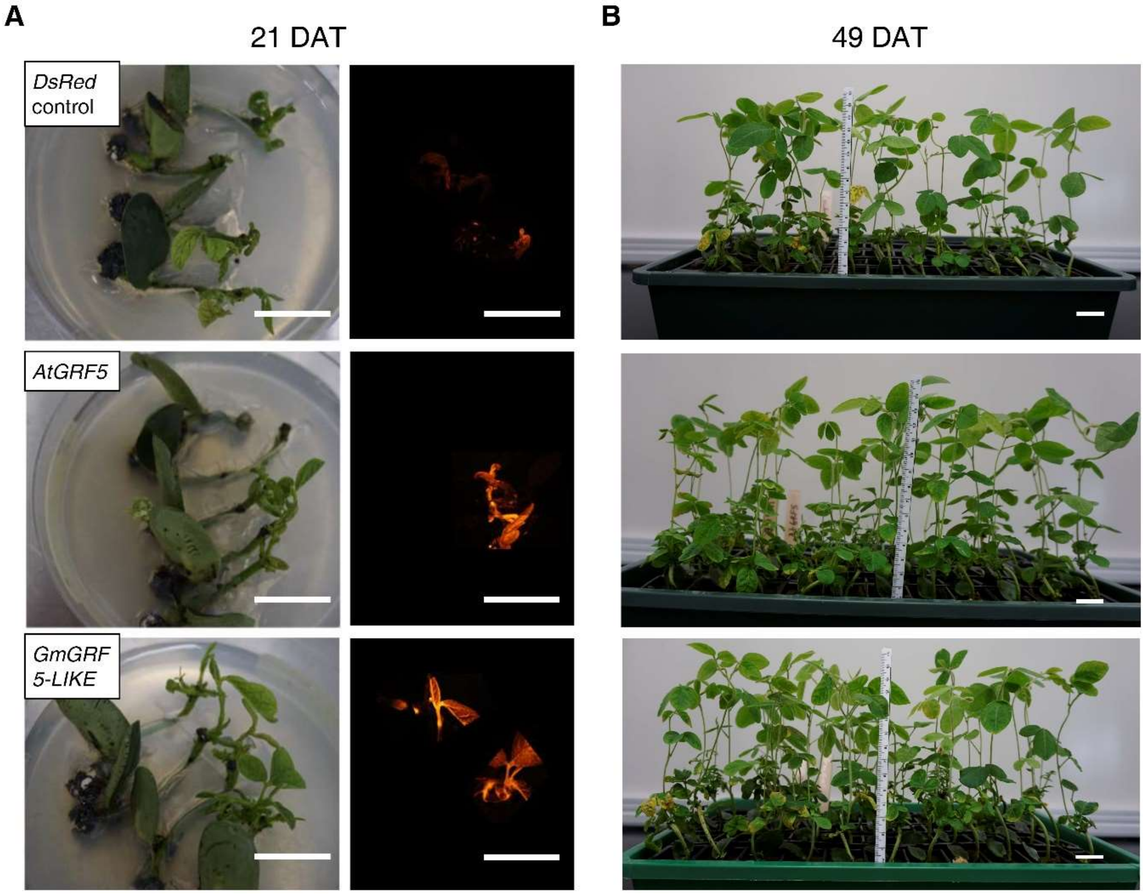


**Supplementary Figure 4**. Shoot development and DsRed fluorescence on soybean (cv. Jake) explants transformed with *DsRed* control, *AtGRF5*, or *GmGRF5-LIKE* using the primary-node method. **(A)** Representative pictures from one repetition of explants transformed with *DsRed* control, *AtGRF5*, and *GmGRF5-LIKE* showing shoot development at the primary-node 21 days after transformation (DAT) under bright field (left) *DsRed* fluorescence of shoots (right). **(B)** Continued development of shoots 49 DAT. Note: *DsRed* pictures are a composite and positioned in approximate position for orientation purposes. Scale bar = 2 cm.

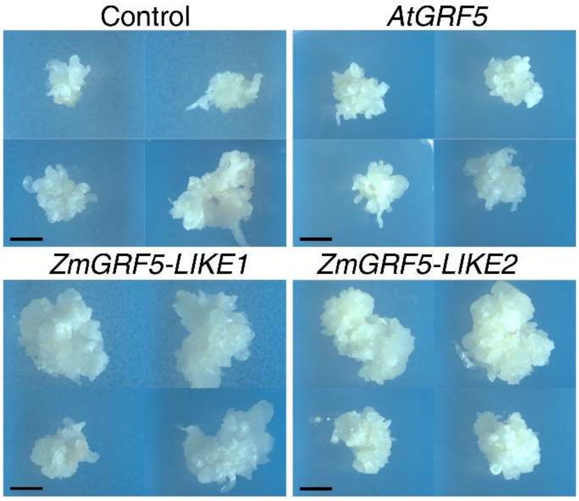


**Supplementary Figure 5**. Representative embryogenic calluses developed from the transformed scutellum of immature embryos in selection medium, at 39 days after *Agrobacterium* inoculation for the constructs overexpressing tdTomato (Control), *AtGRF5*, *ZmGRF5-LIKE1* or *ZmGRF5-LIKE2*. Scale bar = 3 mm.

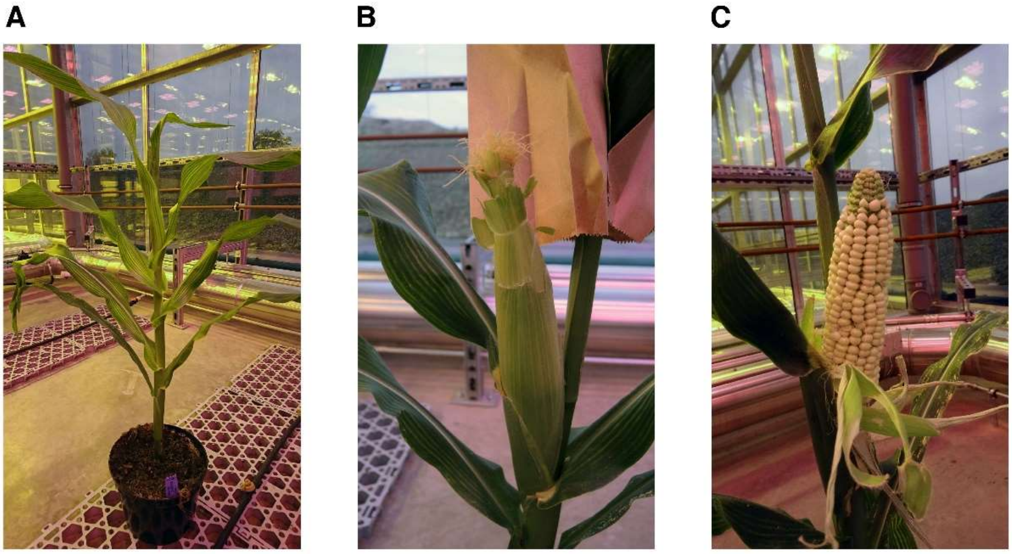


**Supplementary Figure 6**. Analysis of the $T_0$ *ZmGRF5-LIKE* events in maize. Representative pictures at different cultivation steps in the greenhouse are shown. **(A)** Emergence of the tassel in a $T_0$ plant. **(B)** Detail picture of the ear from a self-pollinated $T_0$ plant **(C)** Detail of a $T_0$ ear in maturation.

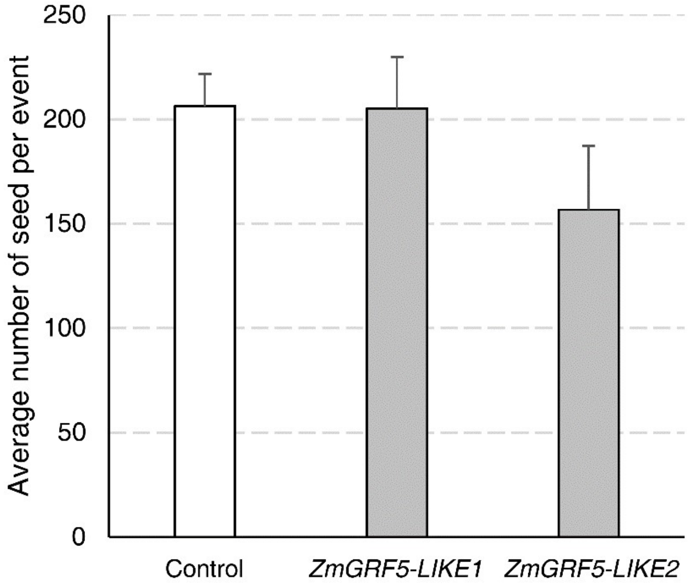


**Supplementary Figure 7**. Seed amount of corn $T_0$ events produced with the *tdTomato* (Control), *ZmGRF5-LIKE1* and *ZmGRF5-LIKE2* constructs. The seed values are indicated as the mean ± SEM from at least 3 independent transgenic events for each construct. Differences in seed number as compared to the events produced with the *tdTomato* construct (Control) are not significant.

| Crop (Transformation method) | *Agrobacterium* strain | Binary vector (Reference) | *GRF* expression cassette | Visual marker | Plant selection |
|---|---|---|---|---|---|
| Sugar beet (A) | *A. tumefaciens* AGL1 | pLH-AB (Hausmann and Töpfer, 1999) | *2x35S*::*AtGRF5*:*NOS* | | *NPTII* |
| | | | *2x35S*::*BvGRF5-LIKE*:*NOS* | | *NPTII* |
| | | | | *2x35S*::*tdTomato*:*pAG7* | *NPTII* |
| Maize (A) | *A. tumefaciens* LBA4404 | pLH-AB (Hausmann and Töpfer, 1999) | *BdEF1*::*AtGRF5*:*ZmEF1* | | *PAT* |
| | | | *BdEF1*::*ZmGRF5-LIKE1*:*ZmEF1* | | *PAT* |
| | | | *BdEF1*::*ZmGRF5-LIKE2*:*ZmEF1* | | *PAT* |
| | | | | *BdEF1*::*tdTomato*:*ZmEF1* | *PAT* |
| Soybean (A) | *A. rhizogenes* SHA017 | pSUN1 (Heim et al., 2007) | *PcUBI4-2*::*GmGRF5-LIKE*:*NOS* | *SuperP*::*DsRed2*:*NOS* | *AtAhasL* |
| | | | | *SuperP*::*DsRed2*:*NOS* | *AtAhasL* |
| Canola (A) | *A. rhizogenes* SHA001 | pSUN1 (Heim et al., 2007) | *PcUBI4-2*::*BnGRF5-LIKE*:*NOS* | *SuperP*::*DsRed2*:*NOS* | *AtAhasL* |
| | | | | *SuperP*::*DsRed2*:*NOS* | *AtAhasL* |
| Sunflower (A) | *A. tumefaciens* EHA105 | pCAMBIA1300 | *35S*::*AtGRF5*:*NOS* | *HaUbi*::*gfp*:*NOS* | *hptII* |
| | | | *35S*::*HaGRF5-LIKE*:*NOS* | *HaUbi*::*gfp*:*NOS* | *hptII* |
| | | | | *HaUbi*::*gfp*:*NOS* | *hptII* |

**Supplementary Table 1**. Detailed description of the *Agrobacterium* strains and constructs used for the transformation experiments for each crop. *Agrobacterium*-mediated transformation method and biolistic transformation are indicated with A and B in brackets, respectively. For sugar beet and corn, the pLH-AB binary vector backbone derived from a nopaline Ti plasmid was used (DCS, DNA Cloning Service e.K., Hamburg, Germany). For soybean and canola the expression cassettes were constructed using a KAN resistant derivative of the dual origin (VS1 and ColE1) binary vector pSUN1. The pCAMBIA1300 binary was used for constructs used in sunflower transformation (CAMBIA, Canberra, Australia). The promoter and terminator for controlling the expression are indicated before and after the GRF5 and the control fluorescent reporter coding sequences in each construct. 2x35S, double cauliflower mosaic virus (CaMV) 35S promoter; NOS, nopaline synthase terminator (Depicker et al., 1982); pAg7, *Agrobacterium*-derived Ag7 terminator; *BdEF1*, *ELONGATION FACTOR-1* promoter from *Brachipodium distachyon*; *ZmEF1*, *Zea mays EF-1* terminator; *PcUBI4-2*, *Petroselinum crispum* ubiquitin gene-derived promoter (Plesch and Ebneth); SuperP, Super promoter (Ni et al., 1995); 35S, CaMV 35S promoter; and HaUbi, promoter derived from the *Heliantus annuus* polyubiquitin gene. As transformation control, the following sequences encoding for fluorescent proteins were used: tdTomato, red fluorescent protein (Shaner et al., 2004); DsRed2, red fluorescent protein derived from *Discosoma* sp. (Invitrogen, Carlsbad CA, USA). The following standard expression cassettes for selection of the transgenic plants were used: neomycin phosphotransferase II (NPTII), phosphinothricin acetyltransferase gene (PAT), acetohydroxyacid synthase gene from *Arabidopsis thaliana* (AtAhasL), hygromicin phosphotransferase II (hptII).

| Crop | Media | Components |
|---|---|---|
| Sugar beet | Callus induction medium | 1X MS basal salts, 15 g/l sucrose and 2 mg/l BAP, 10 g/l agar; pH 5.8. |
| | Co-culture medium | 440 mg/l $CaCl_2x2H_2O$, 170 mg/l $KH_2PO_4$, 1900 mg/l $KNO_3$, 370 mg/l $MgSO_4$, 1650 mg/l $NH_4NO_3$, 20 g/l sucrose, 2 g/l glucose, 0.2 mM acetosyringone and 2 mg/l BAP; pH 5.8. |
| | Shoot regeneration medium | 1X MS basal salts, 30 g/l sucrose, 1 mg/l GA3, 1 mg/l TDZ, 500 mg/l timentin, 100 mg/l paromomycin, 10 g/l agar; pH 5.8. |
| | Shoot multiplication medium | 1X MS basal salts, 30 g/l sucrose, 0.25 mg/l BAP, 100 mg/l kanamycin, 10 g/l agar; pH 5.8 |
| | Rooting | 1X MS basal salts, 0.5 mg/l IBA, 10 g/l agar; pH 5.8. |

**Supplementary Table 2**. Composition of the media used for sugar beet transformation.

| Crop | Media | Components |
|---|---|---|
| Canola | *Agrobacterium* suspension | 1X MS salts and vitamins, 87.6 mM sucrose; pH 5.8 |
| | Germination | 1/2X MS salts and vitamins, 29.2 mM sucrose, 7 g/l Phyto Agar; pH 5.8 in PlantCon™ |
| | Co-cultivation | 1X MS salts and vitamins, 87.6 mM sucrose, 3.1 mM MES, 98.8 mM mannitol, 7 g/l Phyto Agar, 4.5 µM 2,4-D, 500 µM acetosyringone, 1.65 mM L-cysteine; pH 5.6 in plates |
| | Recovery | 1X MS salts and vitamins, 87.6 mM sucrose, 3.1 mM MES, 98.8 mM mannitol, 7 g/l Phyto Agar, 4.5 µM 2,4-D, 300 mg/l timentin; pH 5.6 |
| | Selection | 1X MS salts and vitamins, 87.6 mM sucrose, 2.6 mM MES, 7 g/l Phyto Agar, 0-13.3 µM BAP, 0.54 µM NAA, 0.1-0.29 µM GA, 6 µM $AgNO_3$, 100 nM imazethapyr, 300 mg/l timentin; pH 5.8 |
| | Rooting | 1x MS salts and vitamins, 87.6 mM sucrose, 2.6 mM MES, 7 g/l Phyto Agar, 0.98 µM IBA, 100 nM imazethapyr, 300 mg/l timentin; pH 5.8 |

**Supplementary Table 3**. Composition of the media used for canola transformation.

| Crop | Media | Components |
|---|---|---|
| Soybean Primary-node | *Agrobacterium* suspension | 1/10$^{th}$ B5 salts (G768 Phytotech), 3% sucrose, 20 mM MES, 1X Gamborg's vitamins, 200 µM acetosyringone, 1.44 µM $GA_3$, 5.0 µM Kinetin; pH 5.4 |
| | Germination | 1X B5 salts and vitamins (G398 Phytotech Labs), 2% sucrose, 0.8% Noble agar (A5431 Sigma-Aldrich®); pH 5.8 in PlantCons™ |
| | Co-cultivation | 1/10$^{th}$ B5 salts (G768 Phytotech), 3% sucrose, 20 mM MES, 0.5% Noble agar (A5431 Sigma-Aldrich®), 1X Gamborg's vitamins, 200 µM acetosyringone, 1.44 µM $GA_3$, 5.0 µM kinetin, 4.1 mM L-cysteine, 0.5 mM DTT, 0.5 mM sodium thiosulfate; pH 5.4 |
| | Selection | 1X B5 salts and vitamins (G398 Phytotech Labs), 3% sucrose, 3 mM MES, 1 µM BAP, 5 µM Kinetin, 250 mg/L timentin, 3 µM imazapyr, 0.8% Noble agar (A5431 Sigma-Aldrich®); pH 5.6 in 50 mL dishes |
| | Regeneration | 5.7 µM IAA on Oasis® Wedge® |
| | Rooting | 5.7 µM IAA and 2 µM imazapyr on Oasis® Wedge® |

**Supplementary Table 4**. Composition of the media used for soybean transformation.

| **Crop** | **Assay name** | **Primer or probe name** | **Sequence** |
|---|---|---|---|
| Sugar beet | BvEF1/NPTII | nptII-F | 5′-GCTGTGCTCGACGTTGTCA-3′ |
| | | nptII-R | 5′-CGGCACTTCGCCCAATAG-3′ |
| | | nptII-Probe | 5′-AAGCGGGAAGGGACT-3′ |
| | | BvEF1-F | 5′-CCAAACCTATGGTGGTGGAAACT-3′ |
| | | BvEF1-R | 5′-CTGACGGCAAAACGACCAA-3′ |
| | | BvEF1-Probe | 5′-TCTCAGAGTACCCACCC-3′ |
| Maize | ZmEF1/PAT | Pat-F | 5′-TACGCATACGCCACGCATTA-3′ |
| | | Pat-R | 5′-GGGCGATATACACCGAGTCTTC-3′ |
| | | Pat-Probe | 5′-CGCTCCGCCTACCGT-3′ |
| | | ZmEF1A-F | 5′-GTCCAACAGGGACAGTTCCAA-3′ |
| | | ZmEF1A-R | 5′-CGTCTCCCCCTTCAGGATGT-3′ |
| | | ZmEF1A-Probe | 5′-ACCACCAATCTTG-3′ |
| Soybean | GmLectin/AHAS | Csr1-F | 5’-CCTTGGAGCTATGGGATTTGG-3’ |
| | | Csr1-R | 5’-CCACAACTATCGCATCAGGGTTA-3’ |
| | | Csr1-Probe | 5’-ACAGACGCTCCAATCGCAGCAGGA-3’ |
| | | GmLe1-1664F | 5’-CCTGCAAAGGAGGCTGCTAA-3’ |
| | | GmLe1-1728R | 5’-CATGCGATTCCCCAGGTATG-3’ |
| | | GmLe1-1688Probe | 5’-CCCAAGTCATCATCATGAACCACCCTG-3’ |

**Supplementary Table 5.** Primer and probe sequences used for the TaqMan assays.

| Gene name | Identity |
|---|---|
| *BnGRF5-LIKE* | 84% |
| *GmGRF5-LIKE* | 42% |
| *BvGRF5-LIKE* | 35% |
| *HaGRF5-LIKE* | 34% |
| *ZmGRF5-LIKE1* | 32% |
| *ZmGRF5-LIKE2* | 32% |

**Supplementary Table 6**. Sequence homology of *GRF5-LIKE* genes from different species compared with *AtGRF5*.

| Experiment | Construct | 0-copy | 1-copy | 2-copy | 3+-copy | Total plants |
|---|---|---|---|---|---|---|
| 1 | *DsRed* Control | 1 (6%) | 5 (28%) | 2 (11%) | 10 (55%) | 18 |
| | *AtGRF5* | 0 (0%) | 6 (33%) | 4 (22%) | 8 (44%) | 18 |
| 2 | *DsRed* Control | 0 (0%) | 6 (33%) | 4 (22%) | 8 (44%) | 18 |
| | *GmGRF5-LIKE* | 0 (0%) | 8 (44%) | 6 (33%) | 4 (22%) | 18 |
| 3 | *DsRed* Control | 0 (0%) | 7 (39%) | 4 (22%) | 7 (39%) | 18 |
| | *AtGRF5* | 0 (0%) | 2 (11%) | 9 (50%) | 7 (39%) | 18 |
| | *GmGRF5-LIKE* | 0 (0%) | 7 (39%) | 8 (44%) | 3 (17%) | 18 |
| | Total | 1 (1%) | 41 (33%) | 37 (29%) | 47 (37%) | 126 |

**Supplementary Table 7.** Copy-number analysis confirms transgene integration in $T_0$ soybean plants transformed with *AtGRF5*, *GmGRF5-LIKE*, and *DsRed* vector control using the primary-node transformation method. 126 plants across 3 experiments and 3 repetitions each were sent to the greenhouse and TaqMan assayed for copy number of the *AtAhasL* gene.

| Experiment | Rep | Constructs | Event | $T_0$ copy # | +/- |
|---|---|---|---|---|---|
| 1 | 1 | *DsRed* control | RBMMPW | 1 | 7/5 |
| 1 | 1 | *DsRed* control | RBMMPX | 2 | 5/7 |
| 1 | 2 | *DsRed* control | RBMOAA | 1 | 11/1 |
| 1 | 2 | *DsRed* control | RBMOAK | 1 | 6/6 |
| 1 | 3 | DsRed control | RBMQIU | 1 | 10/2 |
| 1 | 3 | *DsRed* control | RBMQIV | 3 | 12/0 |
| 1 | 1 | *AtGRF5* | RBMNCT | 1 | 6/6 |
| 1 | 1 | *AtGRF5* | RBMNCU | 2 | 10/2 |
| 1 | 2 | *AtGRF5* | RBMNXJ | 1 | 10/2 |
| 1 | 2 | *AtGRF5* | RBMNYE | 1 | 7/5 |
| 1 | 3 | *AtGRF5* | RBMQMG | 1 | 8/4 |
| 1 | 3 | *AtGRF5* | RBMQNJ | 1 | 7/5 |
| 2 | 1 | *DsRed* control | RBMUVL | 1 | 9/3 |
| 2 | 1 | *DsRed* control | RBMUVT | 1 | 8/4 |
| 2 | 2 | *DsRed* control | RBMVSN | 1 | 9/3 |
| 2 | 2 | *DsRed* control | RBMVTX | 1 | 12/0 |
| 2 | 3 | *DsRed* control | RBMXGM | 1 | 11/1 |
| 2 | 3 | *DsRed* control | RBMXQN | 1 | 9/3 |
| 2 | 1 | *GmGRF5-LIKE* | RBMUNB | 1 | 7/5 |
| 2 | 1 | *GmGRF5-LIKE* | RBMUNS | 1 | 9/3 |
| 2 | 2 | *GmGRF5-LIKE* | RBMVSZ | 1 | 9/3 |
| 2 | 2 | *GmGRF5-LIKE* | RBMVTA | 1 | 10/2 |
| 2 | 3 | *GmGRF5-LIKE* | RBMXIL | 1 | 8/4 |
| 2 | 3 | *GmGRF5-LIKE* | RBMXJA | 1 | 11/1 |
| 3 | 1 | *DsRed* control | RBNAXS | 1 | 9/3 |
| 3 | 1 | *DsRed* control | RBNAZC | 1 | 11/1 |
| 3 | 2 | *DsRed* control | RBNCXF | 1 | 6/6 |
| 3 | 2 | *DsRed* control | RBNCYZ | 1 | 11/1 |
| 3 | 3 | *DsRed* control | RBNDWQ | 1 | 10/2 |
| 3 | 3 | *DsRed* control | RBNDXU | 1 | 6/3 |
| 3 | 1 | *AtGRF5* | RBNBBA | 1 | 10/2 |
| 3 | 1 | *AtGRF5* | RBNBAZ | 2 | 11/1 |
| 3 | 2 | *AtGRF5* | RBNDEX | 2 | 8/4 |
| 3 | 2 | *AtGRF5* | RBNDEY | 2 | 11/1 |
| 3 | 3 | *AtGRF5* | RBNDYD | 2 | 11/1 |
| 3 | 3 | *AtGRF5* | RBNDXK | 2 | 10/2 |
| 3 | 1 | *GmGRF5-LIKE* | RBNBBZ | 1 | 12/0 |
| 3 | 1 | *GmGRF5-LIKE* | RBNBGG | 1 | 12/0 |
| 3 | 2 | *GmGRF5-LIKE* | RBNDBQ | 1 | 10/2 |
| 3 | 2 | *GmGRF5-LIKE* | RBNDBR | 1 | 8/4 |
| 3 | 3 | *GmGRF5-LIKE* | RBNDXA | 1 | 10/2 |
| 3 | 3 | *GmGRF5-LIKE* | RBNDXC | 1 | 10/2 |

**Supplementary Table 8.** $T_1$ progeny inherited the transgene from each transgenic event assayed for presence/absence of the *AtAhasL* gene. Twelve immature seeds without seed coat were targeted for sampling from two independent events derived from each experiment, repetition (rep) and construct combination.

| | Observed | | Expected | | | | | |
|---|---|---|---|---|---|---|---|---|
| Progeny name | Transgenic | WT | Transgenic | WT | Degrees of freedom | Chi-square | P-value (two-tailed) | P-value (one-tailed) |
| L1-17 | 22 | 6 | 21.00 | 7.00 | 1 | 0.190 | 0.663 | 0.331 |
| L1-41 | 22 | 8 | 22.50 | 7.50 | 1 | 0.044 | 0.833 | 0.417 |
| L1-43 | 27 | 9 | 27.00 | 9.00 | 1 | 0.000 | 1.000 | 0.500 |
| L1-45 | 29 | 7 | 27.00 | 9.00 | 1 | 0.593 | 0.441 | 0.221 |
| L2-14 | 19 | 8 | 20.25 | 6.75 | 1 | 0.309 | 0.579 | 0.289 |
| L2-16 | 9 | 4 | 9.75 | 3.25 | 1 | 0.231 | 0.631 | 0.315 |
| L2-17 | 17 | 3 | 15.00 | 5.00 | 1 | 1.067 | 0.302 | 0.151 |
| L2-18 | 11 | 2 | 9.75 | 3.25 | 1 | 0.641 | 0.423 | 0.212 |

**Supplementary Table 9**. Transgene segregation analysis in the $T_1$ progeny of 8 independent $T_0$ events in corn. In the progeny name, L1 and L2 corresponds to plants produced by transforming the constructs to overexpress the *ZmGRF5-LIKE1* and *ZmGRF5-LIKE2* orthologs, respectively. Progenies from four $T_0$ events transformed with each *ZmGRF5* ortholog were analyzed. $T_1$ plants were classified as either transgenic or wild-type (WT) based on the qPCR assay. According to the Chi-square test, the difference between observed and expected values was not statistically significant for all analyzed $T_1$ progenies.
